# A GCN1-independent activator of the kinase GCN2

**DOI:** 10.1101/2025.06.29.662197

**Authors:** JiaYi Zhu, Giulia Emanuelli, Glenn R Masson, Vanesa Vinciauskaite, Henriette Willems, Peiqiao Cai, Andrew Lim, Christopher Alan Brown, David Winpenny, Murray Clarke, Rebecca Gilley, Fergus Preston, Jordan CJ Wilson, Aldo Bader, Taufiq Rahman, Joseph E Chambers, John Skidmore, Nicholas W Morrell, Stefan J Marciniak

## Abstract

Mutations of *EIF2AK4*, which encodes the eIF2α kinase GCN2, cause a severe inherited form of pulmonary hypertension called pulmonary veno-occlusive disease (PVOD). Some pathogenic variants of GCN2 are amenable to pharmacological reactivation by low concentrations of ATP-pocket binding inhibitors. Kinase inhibition at modestly elevated concentrations limits the clinical utility of these drugs against PVOD. We therefore performed an *in cellulo* chemical screen for GCN2 activators and identified three structurally distinct compounds with low micromolar stimulatory activities. Unlike previously described GCN2 activators, one of these molecules activated GCN2 independently of GCN1. Modelling supported by structure activity screens suggested it binds within the ATP-pocket of GCN2, but unlike existing ligands does not protrude inward into the allosteric pocket or outward into the solvent. This overcomes a key requirement of other GCN2 activators.

## Introduction

Phosphorylation of eukaryotic translation initiation factor 2 alpha (eIF2α) mediates the response to a variety of stresses by regulating translation, metabolism and cell survival (1, 2). In yeast, GCN2 is the sole eIF2α kinase and triggers the General Control Response to the accumulation of uncharged tRNA and to ribosome stalling (3, 4). Gene duplication gave rise to a family of eIF2α kinases in multicellular organisms linking eIF2α phosphorylation to a variety of other stresses. HRI (EIF2AK1) is activated by haem deficiency, oxidative stress and mitochondrial dysfunction (5–7), PKR (EIF2AK2) responds to cytosolic double stranded RNA during viral infections, while PERK (EIF2AK3) is activated by endoplasmic reticulum (ER) stress (2). Mutations affecting these kinases have been linked to the development of disease and so their manipulation by small molecules is being pursued as a means to develop therapies (2, 8–11).

The eIF2 complex recruits initiator methionyl-tRNA to ribosomes commencing protein synthesis (2). This requires eIF2 to bind GTP, which is subsequently hydrolysed. Bound GTP is replenished by the guanine nucleotide exchange factor (GEF) eIF2B (12), but phosphorylated p-eIF2α binds avidly to eIF2B inhibiting its GEF activity (13). When levels of eIF2-GTP-tRNA-Met subsequently fall, most translation is disfavoured, but some transcripts are translated more efficiently owing to the presence of upstream open reading frames (uORFs) within their 5’ untranslated regions (5’UTRs) (1, 14). ATF4 and CHOP are transcription factors upregulated in this manner during stress to induce genes of the Integrated Stress Response (ISR) (1, 2, 14). Eventually, PPP1R15A (also called GADD34), itself an ISR target, dephosphorylates p-eIF2α to restore the basal state (15, 16).

GCN2 is comprised of five domains—RWD, pseudokinase, kinase, HisRS-like, and C-terminal—and exists constitutively as a homodimer (17, 18). Its kinase activity is stimulated when the RWD domain interacts with colliding ribosomes via the cofactor GCN1 (19–21) or when the HisRS domain interacts with deacylated tRNAs (22, 23). Direct interaction between GCN2 and the ribosome P-stalk can also mediate GCN2 activation during ribosome stalling (24, 25).

Mutations of *EIF2AK4,* which encodes GCN2, cause the aggressive subtypes of familial pulmonary hypertension, pulmonary veno-occlusive disease (PVOD) and pulmonary capillary haemangiomatosis (PCH) (2, 17). Some pathogenic alleles generate hypomorphic variants of GCN2 that we and others have shown can be activated by small molecule inhibitors that bind to the GCN2 ATP pocket, e.g. neratinib and GCN2iB (9, 11, 26). Type 1 inhibitors bind the ATP pocket of the kinase in its active conformation (DFG-Asp-in, αC-helix-in), while type 2 inhibitors bind the inactive conformation (DFG-out, αC-helix-out) (27). Type 1.5 inhibitors, by contrast, bind an intermediate conformation (DFG-Asp-in, αC-helix-out). Several kinase inhibitors classified as type 1 or type 1.5 with respect to their canonical targets can activate GCN2 at low concentrations while inhibiting it at higher concentrations; however, their binding mode to GCN2 remains undefined (9, 11, 26). GCN2iB is a relatively selective type 1.5 inhibitor of GCN2 that activates the kinase at submicromolar concentrations (9, 17). It is hypothesised that drug-induced stabilisation of one GCN2 protomer within a dimer is propagated to the second protomer to cause its activation (11, 28). Several inhibitors of other kinases that are in clinical use or development as anticancer agents activate GCN2 in a GCN1-dependent manner, which may contribute to their therapeutic efficacy (26, 29–31). For this reason, GCN2 activators are being sought as potential anticancer drugs.

The complexity of GCN2 activation, involving direct and indirect detection of amino acid deprivation and ribosome stalling, has hampered efforts to reconstitute GCN2 function *in vitro* (32–34). We therefore performed a cell-based chemical screen for GCN2-dependent activators of the ISR, followed by *in vitro* validation using orthogonal approaches. In doing so, we identified three novel, structurally distinct GCN2 activators. Notably, one of these, designated compound 20, activates GCN2 in a GCN1-independent manner.

## Results

### Chemical screen for GCN2 activators

We generated clonal Chinese Hamster Ovary (CHO) cell lines expressing an ATF4::NanoLuc reporter comprising the 5’ untranslated region (UTR) of human ATF4 mRNA fused to NanoLuc luciferase (Figure 1A) (17). Activation of the ISR could be detected as bioluminescence in response to ISR activators histidinol, tunicamycin and latrunculin A (Figure 1B). Histidinol activates GCN2 by inhibiting histidyl⍰tRNA synthetase, leading to the accumulation of uncharged tRNA^His^. This uncharged tRNA binds to GCN2, relieving its autoinhibition and activating the kinase (35). Latrunculin A sequesters G⍰actin, thereby inhibiting PPP1R15A activity (36, 37). Tunicamycin inhibits protein glycosylation in the endoplasmic reticulum (ER), resulting in activation of PERK (38). A high-throughput screen of the BioAscent library of 123,222 drug-like compounds performed by the ALBORADA Drug Discovery Institute yielded two pools of potential ISR inhibitors and activators. Compounds were classified based on Z-score thresholds derived from the primary screen, with activators defined as those with Z-score > 3. An initial analysis of 121,000 compounds, excluding plates failing quality control, identified 6,521 activators, of which 6,461 overlapped with the subset selected for follow-up screening. Minor discrepancies were restricted to compounds absent from the analysed dataset (e.g. originating from failed plates) or those with Z-scores close to the threshold (3 to 3.02), consistent with limited analytical variation. Using this approach, we assembled a subset of 6,461 compounds enriched for potential ISR activators and screened these in 384-well format at 10µM for 16 hours. To increase sensitivity, screens were performed with test compound alone or combined with the half-maximal effective concentration (EC_50_) of histidinol. This submaximal concentration was used to allow detection of compounds that enhance ISR signalling when GCN2 is partially activated. This yielded 153 compounds that activated the reporter at least three standard deviations above baseline (Z-score > 3) (Figure 1C).

**Figure 1.**
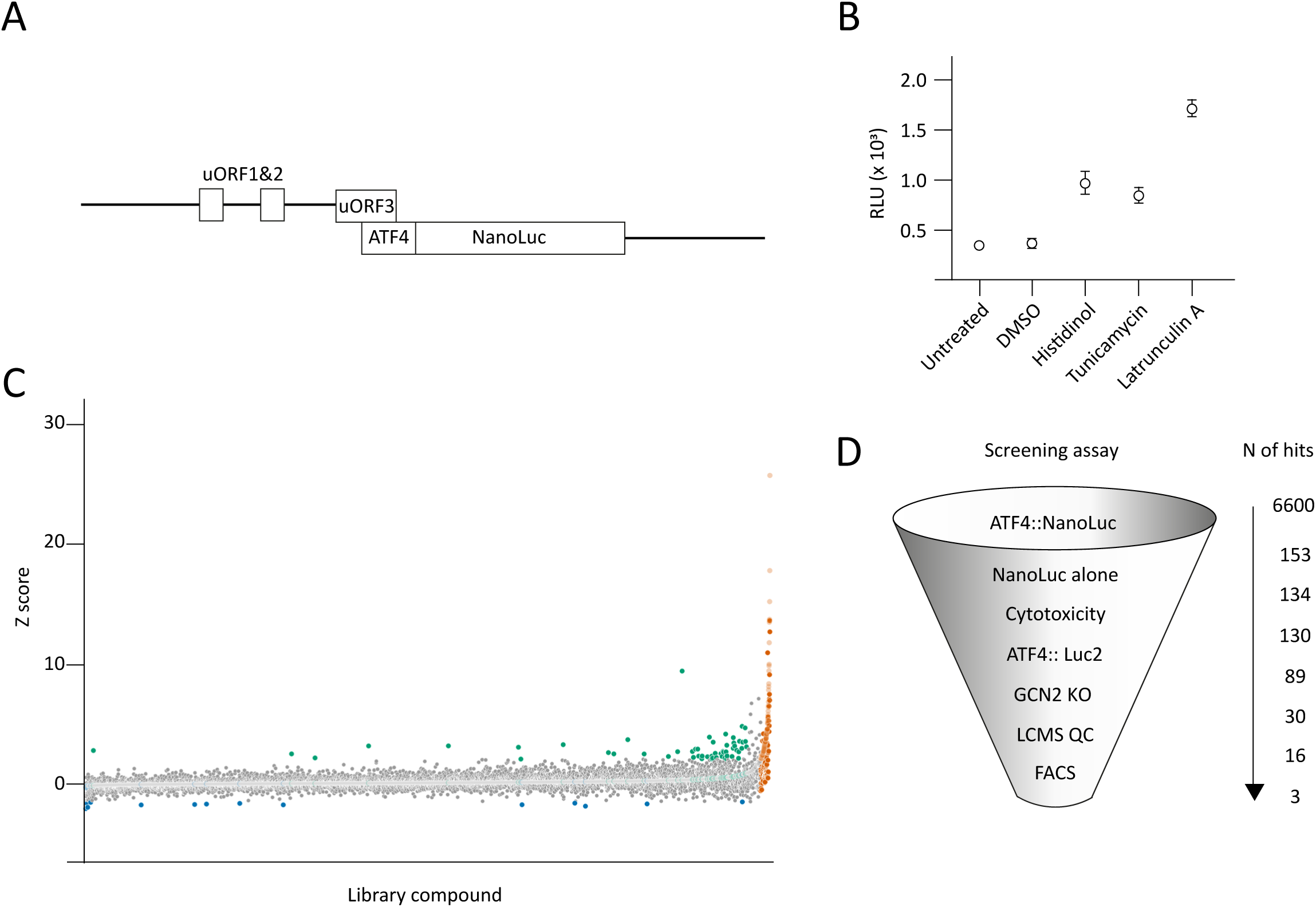
Chemical Screen of novel GCN2-specific ISR-activating molecules. (A) Schematic of ATF4::NanoLuc reporter, exploiting the upstream open reading frames (uORFs). Short uORFs 1, 2 and 3 precede the ATF4 translation start site, with uORF3 overlapping the ATF4 coding sequence, but out-of-frame. During the ISR, skipping of uORF3 leads to increased translation initiation at the ATF4 initiator AUG. (B) Relative Light Unit (RLU) of ATF4 signal in CHO stable cell-line overexpressing ATF4::NanoLuc treated with 1mM histidinol, 2.5 µg/mL tunicamycin, or 1 µM latrunculin A (n=3; ± SEM). (C) Chemical screen of 6,461 small molecules tested with CHO stable cell-line overexpressing ATF4::NanoLuc. The molecules were treated alone or with submaximal ISR activation with 0.3mM histidinol. All molecules were displayed along the x-axis based on their Z-score of relative reporter signal when treated with compound alone (pale symbols). Z-score for treatment with compound and histidinol is also plotted at the same x-axis position (dark symbols). Molecules with Z-score > 3 (i.e., over 3 standard deviations from the mean reporter signal) were identified as hits: ISR activator in orange and ISR augmenter (exaggerating ISR activation by histidinol) in green. (D) Pipeline of molecule short-listing, primary screening using CHO ATF4::NanoLuc reporter (153 hits), NanoLuc alone control (134 hits), selection based on cytotoxicity (130 hits), orthogonal CHO ATF4::FLuc reporter (89 hits), and subsequent GCN2 specificity in GCN2 null reporter cells (30 hits) and compound quality control via LC-MS (16 hits), finally validation by GCN2-specific FACS to a short list of 3 compounds.

The pipeline used to filter hit compounds is illustrated in Figure 1D. Counter screens were performed against constitutively expressed Nanoluc, to exclude trivial activators of luminescent proteins, leaving 134 hits. Cytotoxicity assays excluded immediately toxic compounds, leaving 130. While primary screening was performed in ATF4⍰Nanoluc lines for maximal sensitivity, hits were subsequently re-tested in a second CHO ATF4::luc2 reporter line as a more stringent orthogonal assay to prioritise robust ISR activators. Of the 130 hits identified in the sensitive Nanoluc screen and passing early toxicity assessment, 89 were confirmed in the luc2 assay, consistent with enrichment for higher-amplitude ISR activators under more stringent detection conditions. In subsequent mechanistic studies, the ATF4⍰Nanoluc reporter was again used to take advantage of its higher sensitivity and simpler assay format for multi⍰condition comparisons.

GCN2-deficient ATF4::NanoLuc CHO reporter cells were generated by introducing the reporter into *Eif2ak4-/-* CHO cells (24). Thirty compounds were selected based on GCN2 dependency, where stimulation was lost in GCN2 null cells. Liquid chromatography-mass spectrometry (LC-MS) verified compound purity and quality leaving 16 lead compounds of high confidence from the original library stocks.

### Orthogonal secondary screen

Although hits were prioritised for GCN2 dependence, we performed an additional orthogonal screen to exclude compounds that activate the ISR indirectly via ER stress, which converges on the same downstream outputs. The ISR can be activated in isolation or in combination with other stress signalling pathways, for example as part of the unfolded protein response (UPR) to ER stress (2). To distinguish between inducers of ER stress and selective activators of the ISR, we used dual reporter CHO cells that express a fluorescent ISR reporter (CHOP::GFP) and a reporter of IRE1 activity (XBP1::Turquoise) (39). Histidinol served as a positive control for ISR activation via GCN2, while tunicamycin was used to induce ER stress and activate the UPR. *Eif2ak4^−/-^* CHO cells were used to assess GCN2 dependence, while *Ppp1r15a^−/-^* cells, which are impaired in the dephosphorylation of eIF2α, were used to amplify the effects of ISR activation. Histidinol selectively activated the CHOP::GFP ISR reporter in wildtype and *Ppp1r15a^−/-^*cells, but not in GCN2 deleted cells, validating this system (Figure 2A). Activation of the ISR by tunicamycin was exaggerated in the *Ppp1r15a^−/-^* cells owing to their defective dephosphorylation of eIF2α (wild type vs *Ppp1r15a-/*-, p<0.05). Tunicamycin activated both the CHOP::GFP and XBP1::Turquoise reporter in all three genotypes.

**Figure 2.**
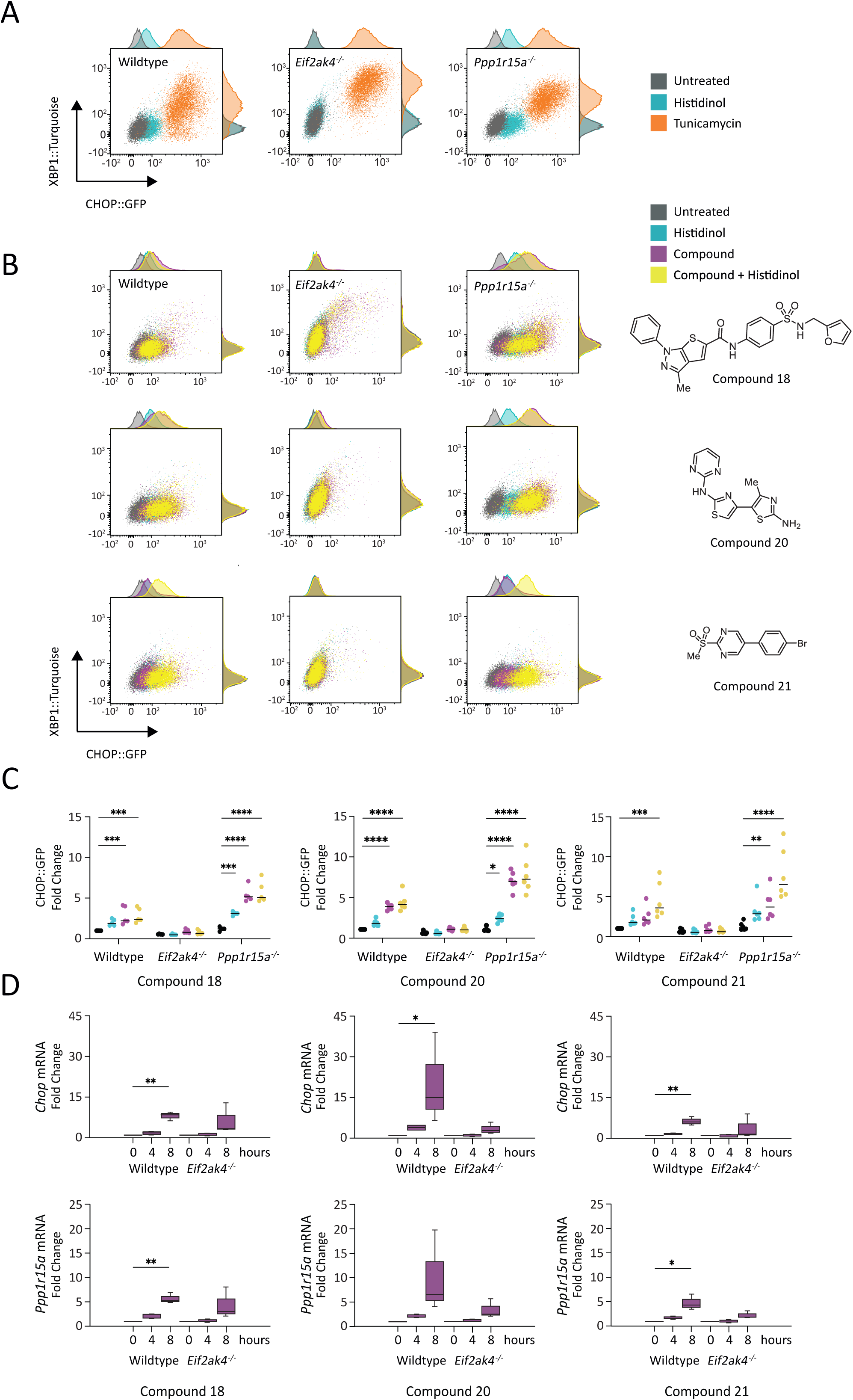
Shortlist and characterisation of three top hits (compound 18, 20, and 21) (A) Representative dot plot of flow cytometry data in CHO CHOP::GFP and XBP1::Turquoise dual-reporter wildtype, *Eif2ak4*Δ, and *Ppp1r15a*Δ cells under 18-hour treatment of 0.75mM histidinol (blue), 2μg/mL tunicamycin (orange), and untreated control (gray) (n=2). Bi-exponential display was used for the plots. The x-axis represents fluorescence signals at 530nm for GFP (CHOP reporter) and y-axis represents fluorescence signals at 450nm for turquoise (XBP1/IRE1 reporter). (B) Representative bi-exponential displayed dot plot of flow cytometry data in CHO CHOP::GFP and XBP1::Turquoise dual-reporter wildtype, *Eif2ak4*Δ, and *Ppp1r15a*Δ cells under 18-hour treatment of 30µM compound 18, 20µM compound 20, and 7.5µM compound 21 alone (magenta) or in combination with 0.75mM histidinol (yellow) (n=6). Histidinol alone is shown in blue and untreated in grey. The x-axis represents fluorescence signals at 530nm for GFP (CHOP reporter) and y-axis represents fluorescence signals at 450nm for turquoise (XBP1/IRE1 reporter). Structures of compounds 18, 20, and 21 shown on the right. (C) Quantification of B (n=6); Two-way ANOVA with Tukey’s multiple comparisons test was performed and the CHOP-GFP signals (530nm) in different treatment groups were compared. *: p ≤ 0.05, **: p ≤ 0.01, ***: p ≤ 0.001, ****: p ≤ 0.0001. (D) Box and whisker plot displaying mRNA fold change of ISR effector CHOP and PPP1R15A after 4- and 8-hour treatment of 30µM compound 18, 10µM compound 20, and 10µM compound 21 as detected by qPCR (n=3). ± SEM, Two-way ANOVA was performed, * p≤0.05, ** p≤0.01.

The 16 lead compounds were next examined both in the presence and absence of a submaximal concentration of histidinol. Using this orthogonal approach, three compounds were found to be GCN2 selective activators of the ISR (Figure 2B&C). Those which induced ER stress or showed minimal GCN2 dependence are not shown. Reporter activation was measured by fluorescence activated cell sorting (FACS) after 18 hours of treatment with 30µM compound 18, 20µM compound 20, and 7.5µM compound 21. The three GCN2 selective activators included compound 18, *N-(4-{[(furan-2-yl)methyl]sulfamoyl}phenyl)-3-methyl-1-phenyl-1H-thieno[2,3-c]pyrazole-5-carboxamide*, containing a furan and two phenyl rings; compound 20, *4’-methyl-N∼2∼-2-pyrimidinyl-4,5’-bi-1,3-thiazole-2,2’-diamine*, containing two aminothiazoles; and compound 21, *5-(4-bromophenyl)-2-(methylsulfonyl)pyrimidine* (Figure 2B).

Larger quantities of compound 18 were then purchased from TimTec, USA (compound ST51216373); compound 20 was synthesised by ChemBridge, USA (compound 5935953); and compound 21 was purchased from Specs, Netherlands (compound AC-907/25005415). After validating their effects using the assays described above, we next confirmed activation of the ISR by these compounds at the transcriptional level at 4 and 8 hours. CHO cells were treated with 30µM of compound 18 or 10µM of compound 20 or 21 for 4 and 8 hours and expression of the ISR target genes *Chop* and *Ppp1r15a* was measured using qPCR (Figure 2D). All three induced ISR gene expression, with compound 20 eliciting the strongest response.

Studies with compound 18 were limited by poor aqueous solubility; therefore, time⍰course analyses focused on compounds 20 and 21. To assess ISR activation over an extended period, live⍰cell luciferase measurements were performed using CHO cells stably expressing an ATF4::Nanoluc-PEST reporter. Both compounds elicited maximal reporter activation between 6 and 8 hours (Figure S1A&B). Compound 20, but not 21, induced a significant GCN2⍰dependent reduction in mRNA translation, as measured by puromycin incorporation, with a progressive effect observed up to 7 hours (Figure S1C-F).

### Cell line differences in GCN2 activator potencies

Neratinib is an ATP-competitive pan-HER kinase inhibitor used clinically in the treatment of breast cancer (40). Recent studies have shown that the sensitivity of glioblastoma cells to neratinib is dependent on GCN2, with GCN2 activation observed at submicromolar concentrations but inhibition at higher concentrations (26). We reproduced this biphasic activation of the ISR reporter in human cervical carcinoma HeLa cells and hamster CHO cells, recording EC_50_ values for activation of 269nM (95% CI 218-322nM) and 690nM (95% CI 634-772nM) respectively (Figure 3A). Disruption of the *EIF2AK4* gene by CRISPR/Cas9 (*EIF2AK4^−/-^*) in HeLa cells abrogated ISR activation confirming the effect to be mediated by GCN2 (Figure 3A, middle panel).

**Figure 3.**
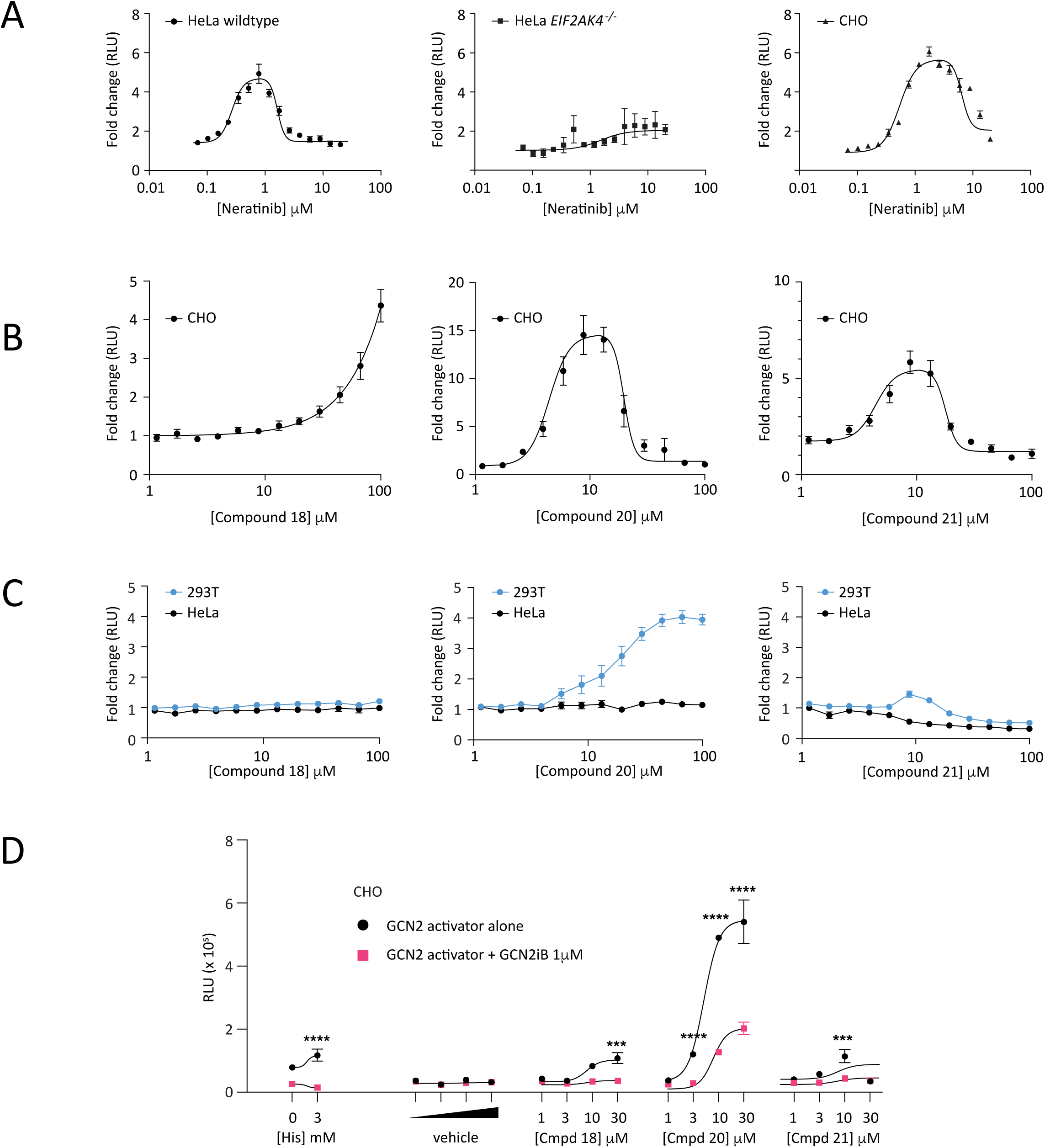
Compound 18, 20, and 21 display cell type-specific differences in ISR activation. (A) Normalised dose-dependent fold change in ATF4 expression by neratinib in HeLa WT, HeLa *EIF2AK4*Δ, and CHO WT ATF4::NanoLuc reporter cells after 16-hour treatment (n=3; mean ± SEM). DMSO was used as vehicle. (B) Normalised dose-dependent fold change in ATF4 expression by compound 18, 20, and 21 in CHO WT ATF4::NanoLuc reporter cells after 16-hour treatment (n=3; mean ± SEM). EC_50_ was estimated using 4-parameter non-linear regression. DMSO treatment was used as vehicle. (C) Normalised dose-dependent fold change in ATF4 expression by compound 18, 20, and 21 in HeLa (black) and 293T (blue) WT cells transiently transfected with ATF4::NanoLuc reporter after 16-hour treatment (n=3, ± SEM). DMSO treatment was used as vehicle. (D) Relative Light Unit (RLU) of of ATF4 signal in CHO WT ATF4::NanoLuc reporter cells treated with 1, 3, 10, 30µM Compound 18, 20, and 21 alone (black) or in combination with 1µM GCN2iB (pink) for 18 hours (n=3; ± SEM). 0.01, 0.03, 0.1, 0.3% DMSO served as vehicle and 3mM histidinol as positive control. Two-way ANOVA was performed *** p≤0.001, **** p≤0.0001.

In CHO cells, compound 18 was the least potent with an EC_50_ of greater than 30µM without obvious inhibition up to 100µM (Figure 3B). Notably, compound 18 failed to activate the ISR measurably in either HeLa cells or human embryonic kidney 293T cells (Figure 3C). Compound 20 activated the ISR in CHO cells in a biphasic manner with an EC_50_ of 4.8µM (95% CI 4.0-5.6µM) (Figure 3B). It was active in 293T cells though with a lower potency, EC_50_ of 17µM, but was inactive in HeLa cells (Figure 3C). Compound 21 also showed biphasic activation of the ISR in CHO cells with an EC_50_ of 5.0µM (95% CI 3.9-6.5µM) (Figure 3B). Weak activation was observed in 293T cells, but none was seen in HeLa cells (Figure 3C). To benchmark ISR activation in these models, each cell type was treated with 3mM histidinol (Figure S2). Reporter activation was most robust in CHO cells, followed by 293T cells, then HeLa cells.

GCN2iB, a potent inhibitor of GCN2, abolished histidinol-induced activation of the ATF4::NanoLuc ISR reporter in CHO cells confirming this to be GCN2 mediated (Figure 3D). Supporting a role for GCN2 in mediating the compounds’ effects, GCN2iB substantially reduced ISR reporter expression by each, though this effect was only partial at concentrations of compound 20 above 10µM (Figure 3D). This might indicate GCN2-independent effects or an ability to partially outcompete GCN2iB for binding at higher concentrations.

We went on to examine downstream cellular consequences of GCN2 activation in multiple models. While compounds did not induce detectable ISR signalling in HCT116 or COS⍰7 cells under the conditions tested, induction of PPP1R15A was observed in Mesobank T12 primary mesothelioma cells, indicating context-dependent biological responses. Moreover, in contrast to GCN2iB (17), the current compounds did not activate disease-associated GCN2 variants linked to PVOD [data not shown].

### Compound 20 activates GCN2 in a GCN1-independent manner

GCN1 mediates the interaction between stalled ribosomes and GCN2 (21, 41). Histidinol is a competitive inhibitor of the histidyl-tRNA synthetase that depletes the cell of charged histidyl-tRNA, thus causing ribosome stalling and activation of the ISR in a GCN1-dependent manner (41–43). Using GCN1 deficient human 293T cells transiently expressing the ATF4::NanoLuc reporter, we confirmed that activation of the ISR by histidinol requires GCN1 (Figure 4A). Low concentrations of GCN2iB (5-70nM) had minimal effect on ISR reporter activation, while neratinib robustly triggered the ISR. The stimulatory effects of neratinib were significantly attenuated in GCN1 deficient cells consistent with the observation that GCN1 knockout renders astrocytoma cells resistant to neratinib (26). By contrast, activation of the ISR by compound 20, chosen for having the largest effect size in 293T cells, did not require GCN1 to activate the ISR reporter (Figure 4A). Indeed, loss of GCN1 modestly increased ISR activation at the highest concentration of compound 20 suggesting a mode of action independent of ribosome stalling. To gain confidence in this observation, we inactivated the *Gcn1* gene in CHO cells by CRISPR/Cas9. Neratinib-induced activation of the ATF4::NanoLuc ISR reporter was lost in GCN1 deficient CHO cells; however, ISR activation by compound 20 was completely insensitive to loss of GCN1 (Figure 4B).

**Figure 4.**
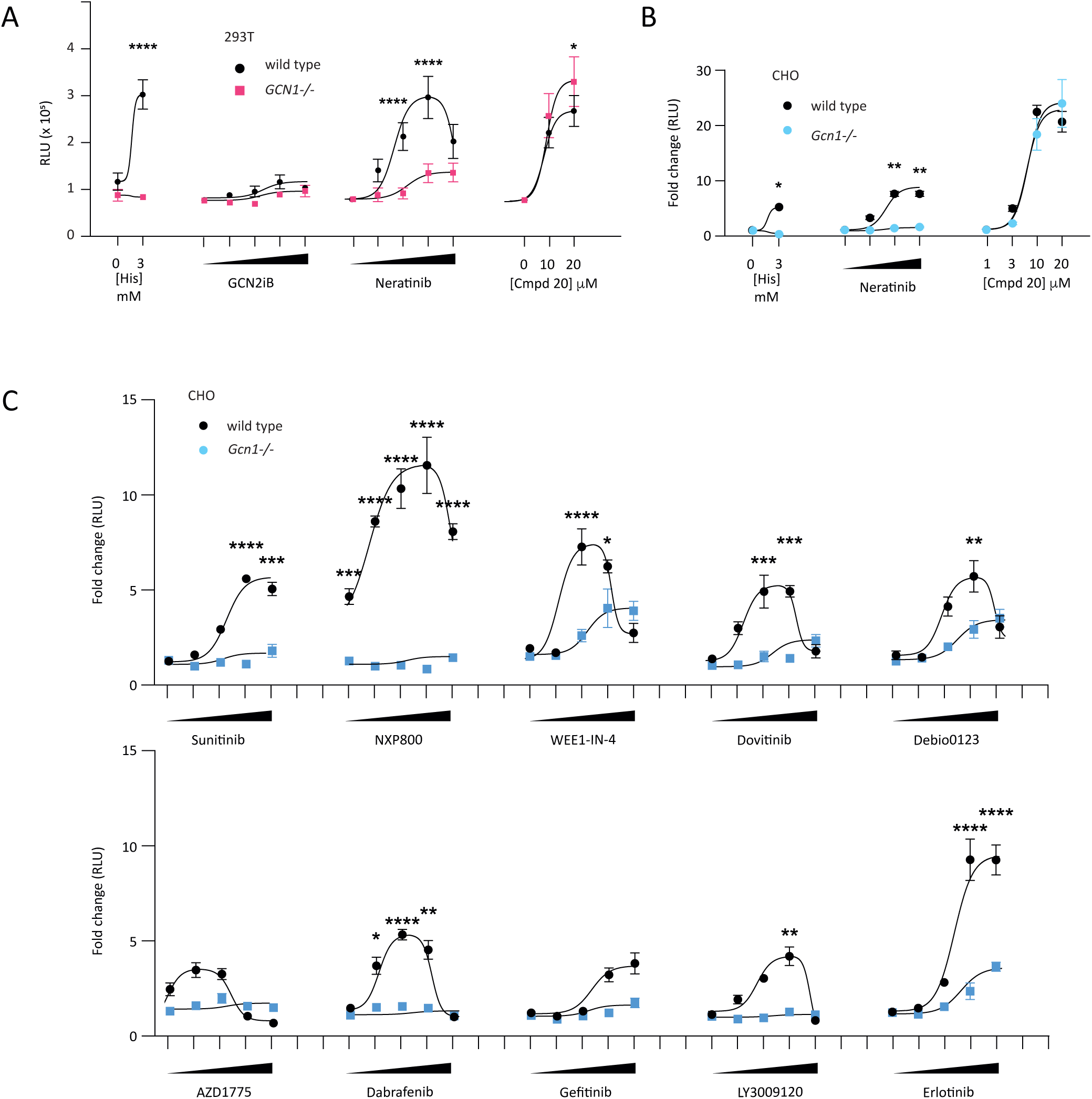
Compound 20 displays GCN1 independence. (A) Relative Light Unit (RLU) of ATF4 signal in 293T WT (black) and GCN1Δ (magenta) cells transiently transfected with ATF4::NanoLuc reporter for 16-hour treatment with compound 20 (10 and 20µM), GCN2iB (5, 10, 35, 75nM), and neratinib (75, 200, 500, 1000nM) (n=3; ± SEM). Two-way ANOVA was performed * p≤0.05, ** p≤0.01, *** p≤0.001, **** p≤0.0001. (B) Normalised fold-change in ATF4 signal in CHO WT (black) and *Gcn1-/-* (blue) ATF4::NanoLuc reporter cells for 18-hour treatment with compound 20 (1, 3, 10 and 30µM) and neratinib (75, 200, 500nM) (n=3; ± SEM). Two-way ANOVA * p≤0.05, ** p≤0.01. (C) Normalised fold-change in ATF4 signal in CHO WT (black) and *Gcn1-/-* (blue) ATF4::NanoLuc reporter cells treated for 19-hours with a panel of ATP-competitive kinase inhibitors reported to activate GCN2 (0.1, 0.3, 1, 3, and 10µM) (n=3-9; mean ± SEM). Two-way ANOVA was performed * p≤0.05, ** p≤0.01, *** p≤0.001, **** p≤0.0001.

No systematic assessment of GCN1 dependence has previously been performed for GCN2 activating drugs. We therefore proceeded to test a panel of ten additional kinase inhibitors that had been reported to activate GCN2 (9, 11, 26). Our panel included examples of multi-targeted tyrosine kinase inhibitors (dovitinib, sunitinib), RAF inhibitors (LY3009120, dabrafenib), inhibitors of the Wee1 kinase involved in G2 cell cycle checkpoint (WEE1-IN-4, AZD1775, Debio0123), an inhibitor of the Heat Shock Factor 1 pathway (NXP800), and EGFR inhibitors (erlotinib and gefinitinib) (9, 11, 26, 31, 44–46). In all cases, a marked reduction in GCN2 activation by the drugs was observed in GCN1 deficient cells (Figure 4C). Most of these molecules failed to activate the ISR in the absence of GCN1 (sunitinib, NXP800, dovitinib, AZD1775, dabrafenib, gefinitinib, LY3009120), while for three drugs weak activation of the ISR was seen at higher concentrations (WEE1-IN-4, Debio 0123, erlotinib). None showed the dramatic GCN1 independence observed for compound 20 in unstressed cells.

To further assess the mechanism of compound-induced ISR activation, we evaluated reporter responses in GCN2-deleted cells (Supplementary Figure S3). Several compounds, including sunitinib, NXP800, WEE1-in-4, Debio0123, gefitinib, and erlotinib, showed reduced reporter activity in GCN2-deficient cells, consistent with GCN2-dependent activation. In contrast, dovitinib, AZD1775 and dabrafenib produced weaker or inconclusive responses even in the paired wild-type lines, limiting analysis. These data allow us to distinguish compounds consistent with direct or GCN2-dependent activation from those more likely to induce ISR indirectly through cellular stress upstream of GCN2. These findings support a distinction between compounds that activate the ISR through GCN2-dependent mechanisms and those that likely act indirectly via alternative stress pathways.

### ATP-competitive interaction with the GCN2 kinase domain

Existing activators of GCN2 have been shown to interact with the ATP binding pocket (9, 11, 26). We therefore used a bioluminescence resonance energy transfer (BRET)-based assay to examine the interaction of the three lead compounds with the kinase domain of GCN2 in cells. Briefly, the N-terminal portion of human GCN2 kinase domain fused to NanoLuc luciferase was expressed in either CHO or 293T cells. These were then treated with an ATP-binding pocked tracer. In this system, energy transfer is proportional to tracer binding to the NanoLuc-tagged kinase domain, and so interactions of other molecules that displace the tracer reduce the BRET ratio (Figure 5A) (47). In both CHO and 293T cells, GCN2iB efficiently suppressed the BRET ratio thus validating the system (Figure 5B). All three lead compounds significantly reduced the BRET ratio, though unlike GCN2iB, compounds 18 and 20 were noticeably more effective in CHO than in 293T cells. Although compound 21 suppressed BRET in both cell types, its effect was less dramatic in CHO cells compared to compounds 18 and 20. The concentrations required to detect target engagement in NanoBRET assays did not directly mirror those required for ISR activation, reflecting the distinction between ligand binding and downstream pathway output.

**Figure 5.**
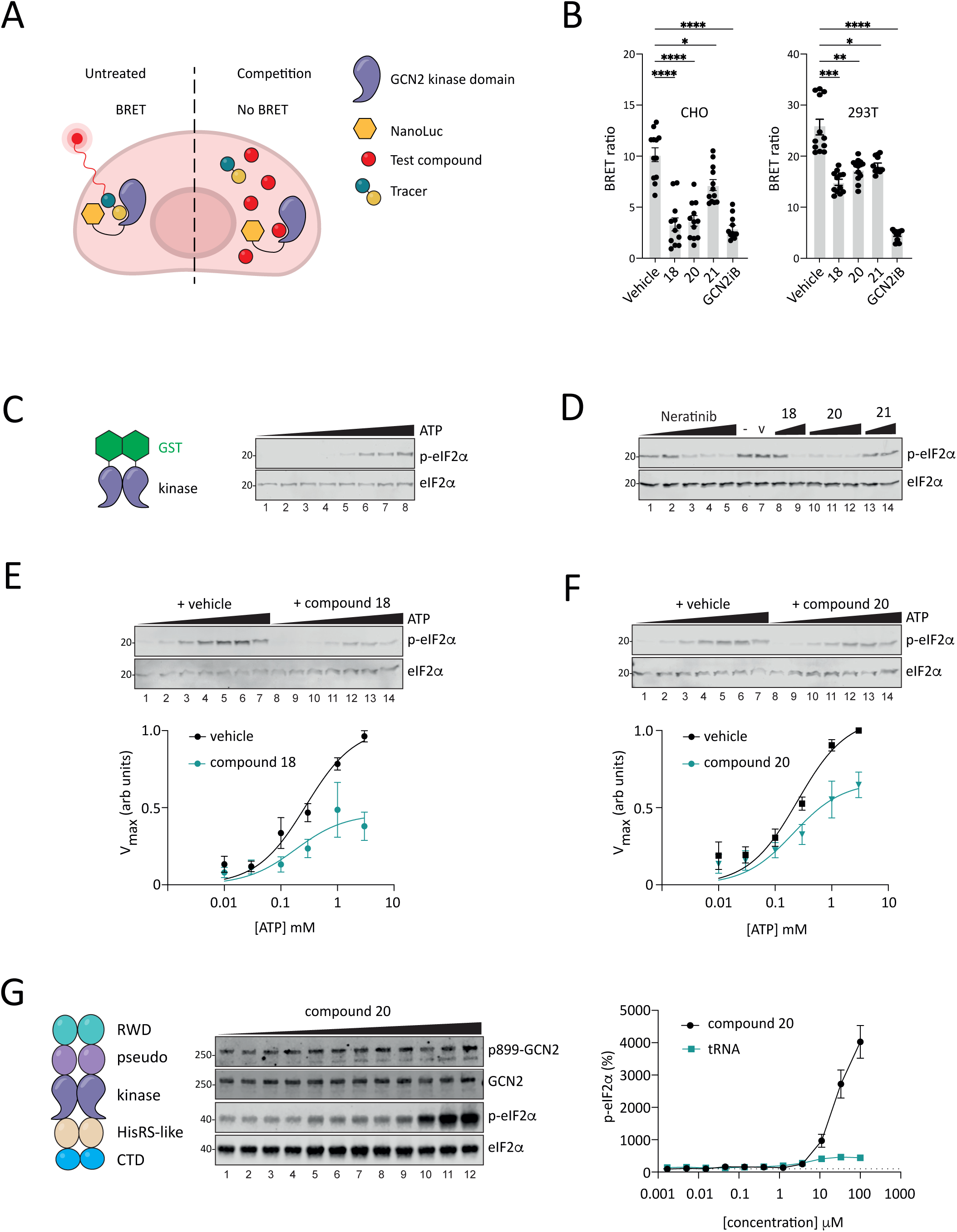
Compound 18 and 20 directly bind to GCN2 in cellulo and in vitro. (A) Illustration of the NanoBRET assay. Left panel displays BRET signal at baseline with tracer binding to GCN2; right panel displays reduced BRET signal due to the addition of a competitive compound that outcompetes the tracer from binding to GCN2. (B) BRET ratio in CHO and 293T WT cells with 2-hour treatment of 3% DMSO, 100µM compound 18, 20, and 21 or 0.5µM GCN2iB. (n=3; ± SEM). Two-way ANOVA was performed. *: p ≤ 0.05, **: p ≤ 0.01, ***: p ≤ 0.001, ****: p ≤ 0.0001. (C) Schematic of GST-GCN2 fusion protein (residues 585–1005) and representative immunoblot of *in vitro* kinase assay. Purified GST-GCN2 was incubated with eIF2α-NTD (residues 2–187) using a ramp of ATP concentrations (0, 0.01, 0.03, 0.1, 0.3, 1, 3, 10mM) (n=3). Total and serine51 phosphorylated eIF2α were detected. (D) Representative immunoblot of *in vitro* kinase assay showing inhibition of eIF2α-NTD (residues 2–187) phosphorylation. Purified GST-GCN2 and eIF2α-NTD was incubated with neratinib (0.075, 0.68, 6.67, 10, 20µM) and compound 18 (100-500µM), compound 20 (100, 200, 500µM), and compound 21 (100-500µM) (n=3). 5% DMSO used as vehicle control. Total and serine51 phosphorylated eIF2α were detected. Molecular size in kDa. (E) Representative immunoblot of *in vitro* kinase assay (n=3) showing inhibition of eIF2α-NTD (residues 2–187) phosphorylation by 300µM compound 18 using a ramp of ATP concentrations (0.01, 0.03, 0.1, 0.3, 1, 3, 10mM); 3% DMSO used as vehicle control. Bottom panel showed Michaelis-Menten kinetics analysis of DMSO control (black) vs compound 18 (green). Molecular size in kDa. (F) Representative immunoblot of *in vitro* kinase assay (n=3) showing inhibition of eIF2α-NTD (residues 2–187) phosphorylation by 100µM compound 20 using a ramp of ATP concentrations (0.01, 0.03, 0.1, 0.3, 1, 3, 10mM); 3% DMSO used as vehicle control. Bottom panel showed Michaelis-Menten kinetics analysis of DMSO control (black) vs compound 20 (green). Molecular size in kDa. (G) Schematic of full-length GCN2 protein and representative immunoblot of *in vitro* kinase assay. Purified full-length GCN2 was incubated with full-length eIF2a using a ramp of compound 20 or tRNA concentrations (0, 0.001, 0.005, 0.01, 0.05, 0.13, 0.4, 1.2, 3.7, 11, 33, 100µM) (n=3). Total and serine51 phosphorylated eIF2a, as well as total and threonine899 phosphorylated GCN2 were detected. On the right showed quantification of serine51 phosphorylated eIF2a with a ramp of tRNA (black) vs compound 20 (green). Molecular size in kDa.

To examine further the interaction of compounds 18 and 20 with GCN2’s kinase domain, we generated in bacteria a recombinant protein comprising Glutathione S-transferases (GST) fused to the kinase domain of human GCN2 (residues 585 to 1005 of NCBI reference Sequence NP_001013725.2). GST was chosen both to facilitate protein purification, and to enhance dimerisation of the kinase domain in order to cause autophosphorylation and constitutive activation. When incubated *in vitro* with increasing concentrations of ATP and a fixed quantity of His6-tagged bacterially expressed eIF2α N-terminal domain (48), GST-GCN2 kinase domain phosphorylated its substrate on serine 51, as detected by immunoblotting (Figure 5C). Neratinib dose-dependently inhibited eIF2α phosphorylation (Figure 5D, lanes 1-5). The lack of activation by neratinib likely reflected the requirement for autoinhibitory domains of GCN2 to observe this effect. Therefore, in this system inhibition of eIF2α kinase activity was used as a surrogate for target engagement. Compounds 18 and 20 also inhibited eIF2α phosphorylation, while compound 21 appeared inactive in this assay (Figure 5D). Compounds 18 and 20 altered the apparent kinetic parameters of GCN2, including an increase in the observed V_max_; however, these data do not allow assignment of a specific inhibition modality (Figure 5E&F).

To test the ability of compound 20 to activate full-length GCN2, we next generated full-length human GCN2 (uniprot ID: Q9P2K8) through baculoviral expression using an N-terminal twin StrepII tag. Despite exerting only a modest effect on GCN2 autophosphorylation at T899 within the activation loop, compound 20 robustly triggered the phosphorylation of full-length eIF2α to an even greater extent than purified tRNAs, a known endogenous stimulus of GCN2 activity (Figure 5G). Of note, uncharged tRNA produced a modest (∼4⍰fold) activation of GCN2 under these conditions, whereas compound 20 induced substantially greater (∼40⍰fold) activation.

### Interaction with other ISR kinases

The eIF2α kinases have a common evolutionary origin and consequently the kinase domains share some structural similarity (49). We therefore examined the effect of the compounds on all four eIF2α kinases *in vitro*. Lower concentrations of ATP (10µM for GCN2, 0.1µM for PERK, 0.15µM for HRI, 0.1µM for PKR) were used to accentuate antagonism at the ATP site. This assay was performed at low ATP concentrations to sensitise detection of ATP-competitive inhibition and was not optimised to detect compound-mediated activation of GCN2. Under these conditions, in contrast to Figure 5G where 100µM ATP was used, activation of GCN2 is not observed with any of the compounds tested. At low ATP, full length, purified GCN2 was efficiently inhibited by GCN2iB with an IC_50_ on 1.55nM. None of the screen hits reached 50% inhibition at the highest concentration tested (3µM) though compound 20 and to a lesser extend 18, appear to cause weak inhibition (Figure 6A). No inhibition was observed with the test compounds again purified PERK (Figure 6B). The PERK inhibitor GSK2606414 served as a positive control with an expected IC_50_ of 0.66nM. Weak inhibition of HRI by compounds 18 and 20 was observed but failed to reach 50% (Figure 6C). In this system, GCN2iB was found to inhibit purified HRI with an IC_50_ of 2.76nM, similar to that observed against GCN2. By contrast, both compounds 18 and 20 inhibited PKR *in vitro*: compound 18 with an IC_50_ of 4.50µM and compound 20 with an IC_50_ of 3.75µM (Figure 6B-D). GCN2iB inhibited PKR with an IC_50_ of 3.26µM. Compound 21 was inactive against each of the four eIF2α kinases in this assay.

**Figure 6.**
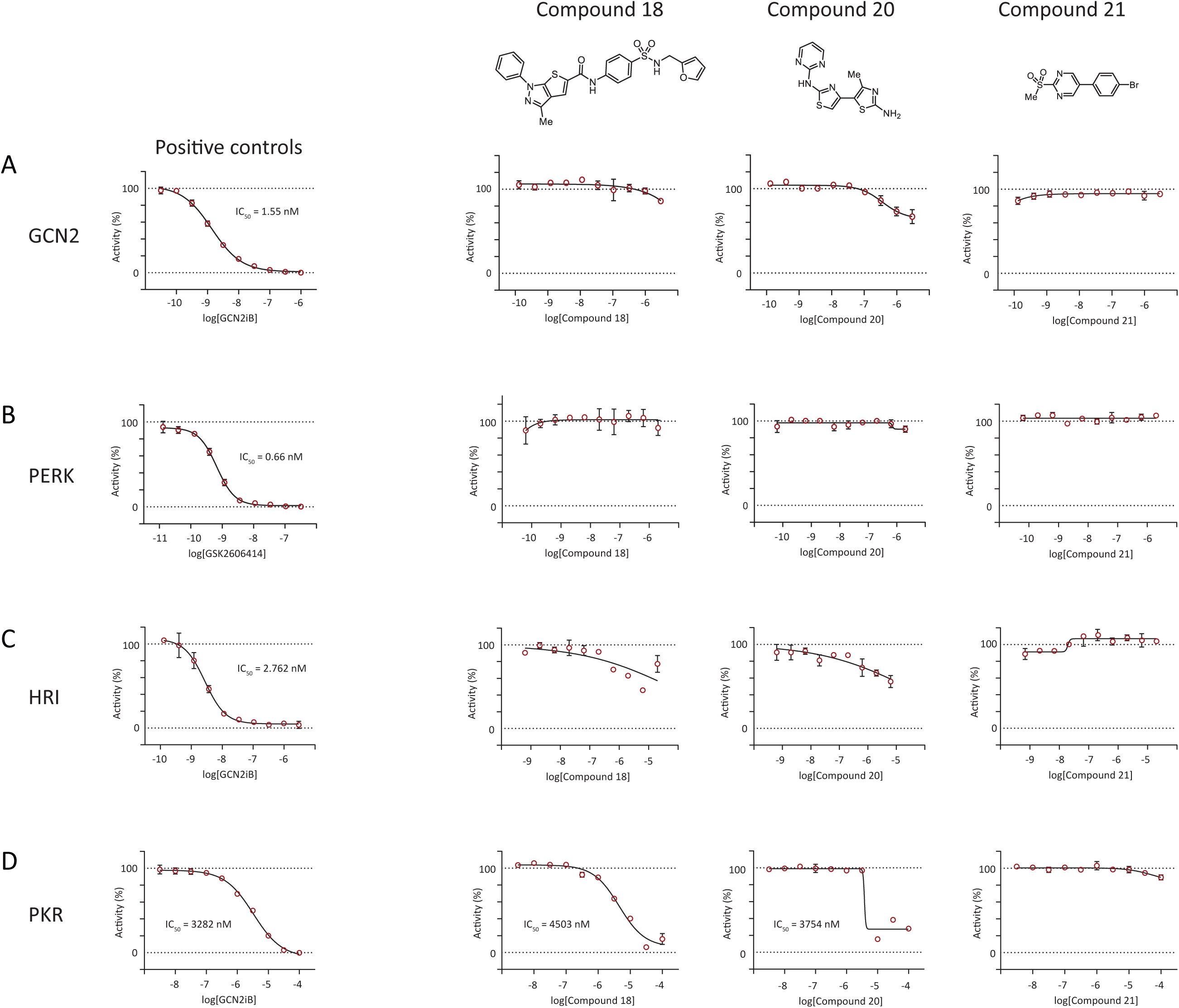
Compound activity across eIF2α kinases using *LanthaScreen in vitro kinase activity assay*. Left panels: dose-dependent inhibition of GCN2, PERK, HRI or PKR with positive control drug GCN2iB or GSK2606414. Right panels: effect of concentration ranges of compounds 18, 20, and 21 in this system revealed weak inhibition of GCN2, HRI, and PKR. Low ATP concentrations were used to accentuate inhibition, and this masks the stimulatory effects on GCN2.

### In silico docking of compounds 18 and 20 to the GCN2 kinase domain and structure-activity relationship (SAR) analysis

We used the SiteMap module from Schrodinger Suite to identify binding sites based on drug interactions with nearby GCN2 residues. Published crystals structures of the GCN2 kinase domain (see methods) show the ATP binding pocket comprises a large solvent exposed region and a buried hydrophobic region that extends beyond the gatekeeper residue (Met802) and towards the ⍰C-helix (Figure 7). Compound 18 was predicted to occupy a similar pocket to that of the ligand in the PDB 6N3N crystal structure demonstrating a Type 1.5 binding configuration, characterised by the DFG loop being “in” and the ⍰C-helix being in the “out” conformation (10). The furan motif of compound 18 was predicted to be buried in the allosteric pocket around αC-helix, between gatekeeper Met802 and Leu640 (Figure 7D). The compound’s sulfone was predicted to interact with Phe867 of the conserved DFG-motif. In neratinib, a triad of hydrogen bonding was predicted between the 8-position of the isoquinoline core (Ar-H bond), the amine linker (H bond) and the 2-position of the fluorobenzene ring (Ar-H Bond) with Asp866, similar to the predicted interaction between the amide linkage of compound 18 and Asp866 (Figure 7B). Both neratinib and compound 18 were also predicted to protrude into the solvent exposed space (Figure 7B-E). However, the interaction of neratinib with Cys805 of the hinge region through the nitrile of isoquinoline core was not present in compound 18.

**Figure 7.**
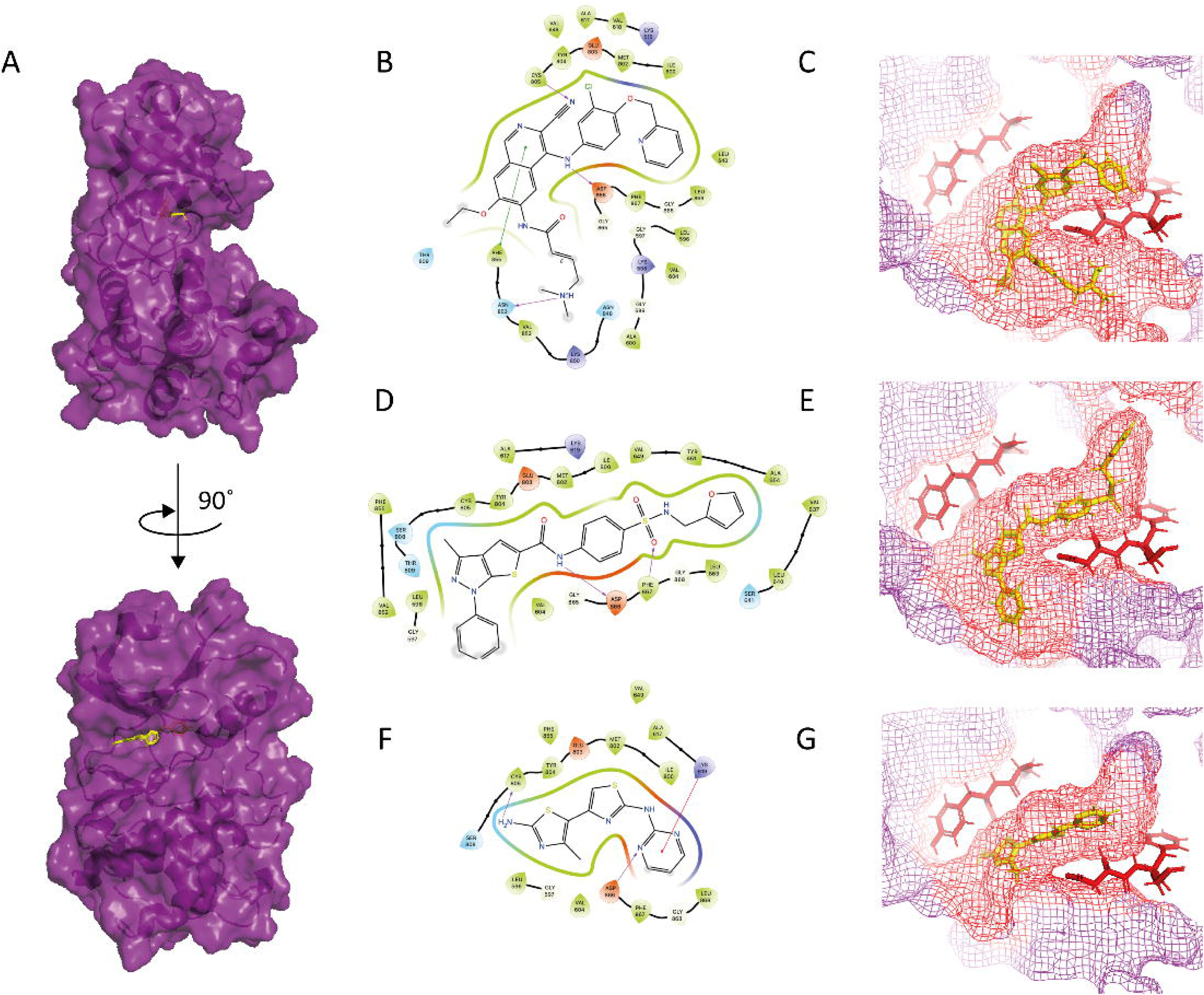
Structural modelling. (A) Illustration of GCN2 kinase domain from up (N-terminus) to down (C-terminus) with illustrative ligand (yellow) in ATP-binding pocket. (B) 2D ligand interaction diagram of neratinib, (D) compound 18, and (F) compound 20. Violet, green, and red colored lining indicate hydrogen bond, π-π, and cation-π interactions, respectively. Polar and hydrophobic residues are highlighted in blue and green, respectively. Mesh representation of the ATP-binding pocket in GCN2 kinase domain was shown with ligand (yellow): (C) neratinib, (E) compound 18, and (G) compound 20. Key residues surrounding the ATP-binding pocket, including the DFG motif (D866-G868) and EYC residues (E803-C805) following the gatekeeper M802 were labelled red. Figures adapted from crystal structure 6N3N.

Due to the metabolic liabilities of furans in medicinal chemistry programs, we sought to explore the importance of this motif for binding and activity using available structural analogues. Replacement of the sulfonamide furan motif with some benzene derivatives including phenolic (BCC0106222), 2-bromobenzene (BCC0113320), 3-acetylbezene (BCC0104214), preserved ISR activation while others (BCC0073242 and BCC0043997) resulted in a loss of activity (Supplementary Table 1). Incorporation of heterocyclic ring systems (BCC0073142, Z56783036) or modification of solvent exposed region (Z88478561, Z226209130, Z11741677) were similarly not tolerated (Supplementary Table 1). However, modifying the furan motif to benzene derivative 2-bromobenzene (BCC0113320) and heterocyclic ring (BCC0077119) preserved GCN2-specific ISR activation.

Unlike compound 18 and neratinib, compound 20 was not predicted to protrude into the allosteric pocket (Figure 7F&G). Analogues extending further towards the ⍰C-helix and protruding into the solvent-exposed region were inactive (e.g., Z118606822), suggesting that the small size of compound 20 may be advantageous, fitting into the ATP-binding pocket of GCN2 and facilitating stronger activation (Supplementary Table 1). Hydrogen bond interaction was predicted between the NH_2_ of the compound 20 aminothiazole with the carbonyl of the hinge region Cys805. Additional interaction was predicted between the linker NH of compound 20 and backbone carbonyl of Asp866 in the DFG-motif. Such 2-aminopyrimidine rings are present in many kinase ligands (50). Rather than this interacting with hinge region residues, this moiety in compound 20 was predicted to form a pi-cation interaction with Lys619 and a hydrogen bond with DFG Asp866, as has been reported for other ligands of kinase ATP pockets (10). Replacement of this pyrimidine ring with 2-pyridine analogues (BCC0084738, BCC0079350 and BCC0083923) was well tolerated, likely due to the conserved hydrogen bonding interaction between the pyridine nitrogen and Asp866. Removal of a nitrogen from the ring system, such as BCC0057014, BCC0089519, BCC0084744, and BCC0084833 all resulted in a loss of activity, possibly due to their lack of interaction with Asp866 (Supplementary Table 1). Due to toxicity issues surrounding 3-aminothiazoles, modifications were made to replace the terminal thiazole ring system with phenyl derivatives (BCC0080152, BCC0075509, BCC0084830, BCC0057431 and BCC0080200). However, these resulted in a decrease in activity towards GCN2, suggesting that there might be a steric component contributing to activity. Replacing the thiazole with a minimally steric affecting thiophene (BCC0079876) also led to activity loss, possibly due to the twisting of central thiazole ring that impaired hydrogen bonding between pyrimidine nitrogen and Asp866. We were therefore unable to improve upon compound 20’s activity but gained valuable SAR information for future development.

To provide additional context on the developability of the identified compounds, we performed an *in silico* assessment of key physicochemical and pharmacokinetic properties. Predicted blood–brain barrier (BBB) permeability scores suggested a gradient across compounds, with compound 18 scoring 5.13 (indicative of a high likelihood of central nervous system penetration), compound 20 scoring 4.29 (moderate predicted penetration), and compound 21 scoring 3.35 (lower likelihood of brain exposure), consistent with its greater polarity. Analysis of lipophilicity, reflecting how readily a compound partitions into lipid membranes, showed that compound 18 has relatively balanced properties (cLogP 1.84; logD 1.98) but very low aqueous solubility (0.07), which may limit bioavailability. Compound 20 exhibited more moderate lipophilicity alongside improved solubility (0.67), whereas compound 21 was the most lipophilic and the least soluble, indicating a less favourable balance of properties. As expected, solubility decreased with increasing molecular weight and lipophilicity. Taken together, these data indicate that compound 20 achieves the most favourable balance between solubility, lipophilicity, and predicted tissue distribution, consistent with a more drug-like profile and supporting its prioritisation for further development.

## Discussion

The ISR plays a critical role for maintaining homeostasis under diverse stress conditions and its disruption results in a variety of disease states (2). The observation that some pathogenic variants of GCN2 are amenable to pharmacological activation led us to seek novel GCN2 activators. Our chemical screen identified three chemically distinct GCN2 activating molecules, one of which is, to our knowledge, the first to show independence from GCN1 for its activity. This may prove useful as a tool compound for the study of GCN2 in cells and using purified components since it overcomes at least one hurdle to kinase activation.

Recent work suggests that the ATP-competitive modulator GCN2iB can activate GCN2 independently of GCN1 under specific conditions using a GCN2 E26A mutant (9). In our hands, we did not observe GCN1⍰independent activation with GCN2iB at the concentrations tested. This discrepancy may reflect a narrow concentration window for GCN1⍰independent activation or context⍰dependent effects of the E26A mutation. These findings raise the possibility that GCN1⍰independent activation of GCN2 may occur under specific conditions or with distinct classes of compounds.

Compounds 18 and 20 demonstrate weak inhibition of the related kinase PKR. Previous structural analyses have shown the kinase domains of PKR and GCN2 to be highly conserved, particularly at the ATP-binding pocket, which likely contributes to this effect. Indeed, a previous publication reported that other small molecule inhibitors of GCN2, including a series based on a triazolo[4,5-d]pyrimidine scaffold, demonstrated cross-inhibition of PKR (51). Furthermore, the PKR inhibitor C16 has been shown to activate GCN2 *in vitro* (11). Intriguingly, viral infections or synthetic dsRNA known to activate PKR, have also been shown to trigger GCN2 in some circumstances, though if this relates to structural similarities of their kinase domains has not been studied (11, 52, 53). While the functional relevance of PKR inhibition in our cellular systems is uncertain, these observations highlight the potential for kinase cross-reactivity, which will be important to address in future studies.

A remarkable finding in the current study was the lack of dependence on GCN1 for compound 20 to activate GCN2 in cells. During amino acid starvation, ribosome-associated GCN1 facilitates binding of uncharged tRNAs to GCN2, thus facilitating GCN2 activation (21, 54, 55). GCN1 also functions as a disome detector, enabling GCN2 activation in response to ribosome collisions (41). In genetic screens identifying modifiers of GCN2 activation by a small molecule, such as the dual HER2/EGFR inhibitor neratinib and the WEE1 inhibitor AZD1775, GCN1 has repeatedly been found to be necessary (26, 30, 31). We were able to confirm this GCN1 dependence for neratinib in the present study. It was therefore surprising that GCN2 activation by compound 20 did not appear to require GCN1 in human or hamster cells. While GCN1 is essential for the activation of GCN2 by amino acid starvation in yeast (56, 57), *in vitro* studies have shown that GCN2 activation by the ribosome P-stalk or by deacylated tRNAs does not absolutely require GCN1 (24, 25). Yet we found compound 20 able to activate purified full-length human GCN2 to a far higher level than was possible with purified tRNAs. The markedly greater activation observed with compound 20 compared with uncharged tRNA *in vitro* raises the possibility that such compounds may partially bypass regulatory constraints on GCN2 activation, including those normally mediated by GCN1. Our modelling suggests that unlike neratinib, compound 20 can fit within the ATP-pocket without needing to protrude into the solvent, nor extending deeply into the hydrophobic region beyond the gatekeeper Met802 residue towards the ⍰C-helix. Indeed, analogues extending in these directions were inactive, perhaps suggesting value to this molecule’s small size. This may prove useful as a tool compound for studies in which GCN1 is unavailable or non-functional.

There are limitations to our study. So far, our experiments have been limited to cultured cells, purified proteins, and *in silico* modelling. We have yet to test our compounds in an animal model. Although our pipeline excluded molecules showing toxicity, it is likely that prolonged activation of the ISR will have toxic consequences via targets including CHOP and PPP1R15A (16). Our most promising molecule, compound 20, contains a 2-aminothiazole. Although known to be a versatile scaffold, appearing in several clinically useful drugs including anti-inflammatory, anti-microbial and anticancer agents, 2-aminothiazole is sometimes considered a toxicophore owing to toxic consequences following its metabolism, and as a pan-assay interference compound (PAINS), owing to frequent false positives in high-throughput drug screens (58–60). Our use of orthogonal approaches to support its activity as a GCN2 activator, mitigate against it being a non-specific hit. Nevertheless, if this compound were to be developed further, attempts to ‘design out’ this feature by, for example, replacing the sulphur for an oxygen (thiazole to oxazole switch) could be attempted. Indeed, computational modelling reveals an adjacent NH which should form a stronger H-bond to oxygen of an oxazole, which may result in an improved potency to GCN2. Another potential limitation of our study is the use of existing analogues in SAR studies, the so-called ‘analogue by catalogue’ approach. Although less systematic than chemically synthesising further series *de novo*, it was cost effective and generated useful information on the tolerability of modifications of our lead scaffolds. Finally, our compounds were identified in a screen of CHO cells and appear to be more effective in these cells that in HeLa and 293T models. The absence of detectable activity in HeLa cells does not appear to be a species effect, since some activity was observed in human 293T cells and when using recombinant human GCN2 protein. It is likely therefore to reflect cell-type effects, since cancer cell lines are known to demonstrate dramatic differences in drug efficiency, even when using different strains of the same cell line (61). Moreover, histidinol-induced ISR activation showed a clear hierarchy across cell lines, with CHO cells being the most responsive and HeLa cells the least. We also evaluated the reported GCN2 activator HC⍰7366 in our CHO ATF4::NanoLuc reporter system. In this context, HC⍰7366 showed limited activity relative to compound 20, despite its reported efficacy in other cellular systems and *in vivo* (data not shown). This highlights potential context dependence in small⍰molecule activation of GCN2 and limits direct cross-study comparison. This should be considered carefully when using these compounds as tools or for further drug development.

In summary, we performed a chemical screen for novel activators of GCN2. We identified three chemically distinct scaffolds, and demonstrated one to be independent of the GCN2 cofactor GCN1.

## Methods

### Screen molecules

The initial screen used compounds sourced from the BioAscent library and all molecules dissolved in DMSO. Some follow-on compounds for SAR were sourced from Enamine. Subsequently, compound 18 was purchased from TimTec, USA (compound ST51216373). Compound 20 was synthesised by ChemBridge, USA (compound 5935953). Compound 21 was ordered from Specs, Netherlands (compound AC-907/25005415).

### NanoGlo® luciferase assay

pGL4.2 ATF4-Nluc and pGL4.2 ATF4-Nluc-PEST® were described previously (17). In 384-well white microplate, test compounds, 3mM histidinol or vehicle control were plated as 5x solution in a volume of 5μL in 10% FBS-Nutrient Mixture F-12 Ham supplemented with 1x GlutaMAX™. CHO wildtype and *Eif2ak4-/-* ATF4::NanoLuc reporter cells were subsequently seeded at a density of 1.5×10^3^ cells per well in 20μL. After 16 hours at 37°C, NanoGlo® luciferase assay reagent was added to each well and mixed with orbital shaking for 1 minute. After 10 minutes of incubation at room temperature, bioluminescence signal (360-545 nm) was acquired on a Tecan Spark plate reader. Cells carrying a ATF4::nanoLuc-PEST reporter were used for live-cell assay, allowing detection of time-sensitive ATF4 signals (17). Cells (1.25×10^4^ cells per well) were plated overnight on white, clear bottom 96-well plate. Next day, 1μL 100x Nano-Glo® Endurazine™ substrate was mixed with cell culture medium and added to cells and incubated for 2 hours to reach stabilisation. Compounds were added to the wells and luminescence signals were measured using a Tecan CM Spark plate reader at 37°C in a humidity cassette for 48 hours. Data analysed using Graphpad Prism 10.

### Bioluminescence Resonance Energy Transfer (BRET)

NanoBRET assay was performed according to user’s manual. Briefly, 4×10^5^ cells/mL (HEK293T or CHO) were transfected with 100ng of GCN2 kinase domain::nanoLuc reporter and 900ng of transfection carrier DNA mixed in 100μL Opti-mem (no phenol red) using lipofectamine LTX, and plated in 384-well cell-culture white microplates (40μL/well, 8×10^3^ cells). Twenty-two hours after transfection, tracer was added to each well and mixed on an orbital shaker. Test compound (final concentration 10, 30, 100μM), DMSO control (final concentration 0.1, 0.3, 1%), and positive control GCN2iB (final concentration 1μM) were added to each well and mixed. After 2 hours of incubation, NanoBRET™ Nano-Glo® substrate-extracellular NanoLuc® Inhibitor reagent was added to each well before bioluminescence measurement using a Tecan CM Spark plate reader (multi-colour measurement mode: wavelength 1, wavelength 445-470nm, 300ms integration; wavelength 2, wavelength 610-635nm, 300ms integration). Data analysed using Graphpad Prism 10.

### Cell Viability Assay

Compounds were plated into 384-well white microplates and mixed with 20μL of culture medium containing 1.5×10^3^ cells per well, incubated for 16 hours at 37°C. CellTiter-Glo® Assay reagent was added to each well for bioluminescence measurement using a Tecan CM Spark plate reader. Data analysed using Graphpad Prism 10.

### Flow cytometry

CHOP::GFP and XBP1::Turquoise dual-reporter cells were treated with each compound (7.5-20μM) and histidinol (0.75mM) 18-20 hours prior to cell harvesting (39). ER stress was induced using 2μg/mL of tunicamycin. At the time of experiment, cells were trypsinised and collected by centrifugation, resuspended in 4mM EDTA PBS, filtered through a 50μm filter, then analysed using a DB Fortessa flow cytometer. GFP was measured at 530nm and Turquoise was measured at 450nm. Data analysed using FlowJo software.

### Quantitative real time PCR (qPCR)

RNA was reverse transcribed to produce complementary DNA using High-Capacity cDNA Reverse Transcription Kit (Thermo, 4368814) according to the manufacturer’s instructions. To perform real⍰time PCR, the PowerUp SYBR Green Master Mix (Thermo A25742) and the CFX96 BioRad real⍰time PCR machine (Biorad, UK) were used. All samples were measured in duplicate and included water control. Fold changes in expression were calculated using the ΔΔCt method. Hamster qPCR primers include

CHOP (Fw: GGGAGCTGGAAGCCTGGTAT, Rv: GGGACCCCCATTTTCATCTG), PPP1R15A ( Fw: CCTGGTCTGCAAAGTGCTGAT, Rv: CCAGCTCAGTCACTCCCTCTTC), and β-actin (Fw: ACTCCTACGTGGGTGACGAG, Rv: AGGTGTGGTGCCAGATCTTC). Data analysed using Graphpad Prism 10.

### Immunoblotting

Cells were washed with PBS on ice and lysed in low-salt buffer as previously described (17). Following SDS-PAGE (7% gel for GCN2 and 14% gel for eIF2α or alternatively, 5-12% gradient gel), proteins were transferred to nitrocellulose membranes then analysed by immunoblotting. Antibodies used: total GCN2 (Gift from Ron lab: NY168); total GCN2 (1:1000, Invitrogen JA03-83); GCN2-phosphorT899 (1:1000, Abcam, ab75836); β-actin (1:5000, Abcam, ab8226); eIF2α phos-S51 (1:1000, Abcam, ab32157); eIF2α phos-S51 (1:500, Cell Signalling Technology 9721); total eIF2α (1:1000, Invitrogen, AHO0802); total eIF2α (Gift from David Ron lab, CIMR); total eIF2α (1:1000, Santa Cruz Biotechnology sc-133132).

### Recombinant GCN2 kinase domain synthesis and in vitro kinase assay

The human GCN2 kinase domain (residues 585 to 1005) was tagged on its N-terminus with GST by cloning into the vector pGV67 by Gibson assembly. Protein was synthesised in BL21(DE3) E. coli Rosetta cells and purified using glutathione agarose beads. GCN2 kinase domain (0.3μg) was incubated with 100μM ATP, 1μM eIF2α-NTD and test compounds in reaction buffer (50mM HEPES, pH 7.4, 100mM potassium acetate, 5mM magnesium acetate, 250μg/ml BSA, 10mM magnesium chloride, 5mM DTT, 5 mM β-glycerophosphate). Reactions were incubated at 32°C for 10 minutes, then immediately quenched with SDS sample buffer pre-warmed to 95°C. Samples were then analysed by immunoblotting.

### Recombinant full-length GCN2 synthesis and in vitro kinase assay

Expression and purification of recombinant human GCN2 was conducted as described previously (25). Briefly, human GCN2 (uniprot ID: Q9P2K8) was cloned into a baculoviral vector with an N-terminal twin StrepII tag followed by a TEV protease site. GCN2 was then expressed in *Sf9* cells grown at 27 °C for 55 hours. Protein was purified using sequential StrepTrap HP column (GE Healthcare Life Sciences 28-9075-47), HiTrap Q HP column (Cytiva 17115401), Hi Load 16/60 Superdex 200 column (Cytiva/GE Healthcare 28989335). Ten nanomolar GCN2 was incubated at 30 °C for 20 min with 2µM eIF2α and 100µM ATP in increasing concentrations (0.00169-100µM) of deacylated tRNA or compound 20. Samples were then quenched via the addition of SDS-PAGE loading buffer and heating to 95 °C for 3 min, then analysed by SDS-PAGE and immunoblotting: total eIF2α (Santa Cruz Biotechnology sc-133132) 1:1000, phospho-eIF2S1 Ser51 (Cell Signalling Technology 9721) 1:500, total GCN2 (Invitrogen JA03-83) 1:1000, pThr899 GCN2 (Abcam AB75836) 1:1000. Images were captured and quantified in Image Studio Ver 5.2. Data analysed using Graphpad Prism 10.

### Expression and purification of eIF2α

Expression and purification of recombinant human eIF2α was conducted as described previously (25). DNA encoding full-length human eIF2α (NCBI reference number: NP_004085.1) was inserted into the vector pOPTH with an N-terminal His6 tag followed by a TEV protease site. Protein was expressed in BL21 Star (DE3) cells, at 37 °C for 3 hours then purified using a HisTrap HP Column (Cytiva 17524801) as described for GCN2.

### LanthaScreen in vitro kinase activity assay

In 384-well plate format, 1nM human GCN2 (Carna Bioscience, # 05-153) or 1nM human PERK (Carna Bioscience, # 05-155) or 2nM human HRI (Carna Bioscience, # 05-154) or 3nM human PKR (Carna Bioscience, # 05-156) were incubated with 50mM HEPES pH 7.0, MgCl_2_ 10mM, MnCl_2_ 5mM, tRNA 4µM (only for GCN2 assays), ATP (10µM for GCN2, 0.1µM for PERK, 0.15µM for HRI, 0.1µM for PKR), GFP-eIF2α 80nM. Reaction time was 20-30 minutes at room temperature, followed by detection time 50-60 minutes. TR-FRET signal between terbium-labeled antibody and GFP-labeled p-eIF2α read with excitation 340nm and emission of the donor (terbium labeled anti-p-eIF2α antibody: 495 nm) and the acceptor (GFP-labeled p-eIF2α: 520nm) on the Tecan Spark plate reader. IC_50_ values were determined in GraphPad Prism 8.0, using a 4-parameter model: log(inhibitor) vs. response – variable slope.

### In silico modelling

Five reported crystal structures for human GCN2 were considered (PDB 6N3N, 6N3O, 6N3L, 7QWK, and 7QQ6 (10, 62). Of these, 6N3N and 6N3L share ∼100% sequence homology. While 6N3N has unresolved amino acids residues, these are remote from the ATP binding site and should not impact the binding of the ligands. A visual inspection of 6N3O, 6N3N, and 6N3L highlighted that the activation loop in 6N3L is cut short and does not extend beyond the ⍰C-helix, affecting the orientation of the DFG loop and its residues (Asp866, Phe867, Gly868). As such, 6N3L was not considered for further analysis. PDB sequences 7QWK and 7QQ6 share ∼75% sequence homology, with 7QWK containing unresolved residues remote from the ATP binding site. As observed with PDB 6N3L, the activation loop containing the DFG residues is cut short in both cases and does not extend beyond the ⍰C-helix. Inspection of the binding mode of crystallised ligands of 7QQ6 and 7QWK demonstrate that these bind in a Type I binding pocket, compared to Type II binding as observed in 6N3O.

Prior to docking, the protein structure (PDB ID: 6N3N) was put through the Protein Preparation Protocol of Maestro. This protocol added hydrogens, removed co-crystallised water molecules, optimised the receptor’s hydrogen bond network, enumerated bond orders, and performed a restrained minimisation to alleviate backbone clashes with the receptor backbone. The protein was prepared at pH7.4 (63). SiteMap module was used on the prepared protein, with all settings set to default (SiteMap, Schrödinger Release 2024-4, Schrödinger, LLC, New York, http://www.schrodinger.com/). Grid generation was performed using the Glide Docking Protocol in Masestro (Glide, Schrödinger Release 2024-4, Schrödinger, LLC, New York, http://www.schrodinger.com/). This was centred around Met802 and Lys619. High throughput virtual screening (HTVS) setting was used for docking.

---

**Supplementary Figure S1. Kinetics of responses to compounds 20 and 21**

(A-B) Wild-type CHO cells stably expressing the ATF4::nanoLuc-PEST reporter were treated with Nano-Glo and either (A) 13μM compound 20 or (B) 13μM compound 21. Median bioluminescence (fold change normalised to DMSO control) ± 95% confidence. Representative experiment (n=4 technical repeats). (C-F) Representative immunoblot of lysates from wild-type or *Eif2ak4^−/-^*CHO cells treated with 10μM compound 20 or 7.5μM 21 for the indicated times. Immediately before harvesting, cells were treated with 10μg/mL puromycin to label newly synthesised polypeptides. “-” indicates cells not incubated with puromycin. “U” cells were treated with puromycin but without test compound. “CHX” represents the cycloheximide control (100μg/mL). Molecular size in kDa. (E-F) Quantification of puromycinylated proteins normalised to GAPDH. Mean ± SEM. CHO WT (black) and *Eif2ak4^−/-^* cells (turquoise. N = 4 independent experiments. Two-way ANOVA with Šídák’s multiple comparisons test; ***: p ≤ 0.001.

**Supplementary Figure S2. Cell-type differences in response to histidinol**

Fold-change of ATF4::NanoLuc reporter signal in HEK293T, HeLa and CHO cells transiently transfected with reporter and treated for 20 hours with 3mM histidinol. Fold-change calculated relative to vehicle control. Mean ± SEM.

**Supplementary Figure S3. GCN2-dependence of ISR activation by putative GCN2 agonists**

Normalised fold-change in ATF4 signal in CHO WT (purple) and Gcn2-/- (blue) ATF4::NanoLuc reporter cells treated for 19 hours with a panel of ATP-competitive kinase inhibitors reported to activate GCN2 (at 1 and 3µM, sunitinib used at 3 and 10µM, AZD1175 used at 0.3 and 1µM, gefitinib and erlotinib used at 3 and 10µM). DMSO was used as vehicle control. (n=3; mean ± SEM).

## Acknowledgements

SJM was supported by the MRC (G1002610, MR/V028669/1 and MR/R009120/1), EPSRC (EP/R03558X/1), Cambridge Biomedical Research Centre (BRC-1215-20014); British Lung Foundation, Asthma+Lung UK, Royal Papworth Hospital, Alpha1 Foundation. GE was supported by philanthropic funding from Rick Medlock and by awards from EPSRC (EP/R03558X/1), BHF (RE/18/1/34212), Evelyn Trust (Grant 22/03), and the Cambridge-Tsinghua collaborative programme on sustainability and emerging technologies. JZ was supported by the Doctoral Training Programme in Medical Research (DTP-MR; Cambridge Trust). GRM is supported by a Baxter Fellowship from the University of Dundee, and a TENOVUS grant T21/01 119615, and Biotechnology and Biological Sciences Research Council (UKRI-BBSRC) Capital Equipment Grant BB/V019635/1. VV supported by MRC iCase Studentship (MR/R01579/1). This research was supported by the CIMR Flow Cytometry Core Facility. We wish to thank Reiner Schulte and Gabriela Grondys-Kotarba for their advice and support in flow cytometry cell sorting. We wish to thank Heather Harding for helpful discussions and reagents. We wish to thank Steve Andrews, Kushal Rugjee, and other colleagues at the ADDI for help with chemical library preparation. Thanks to Apollo Therapeutics Ltd for conducting the LanthaScreen in vitro kinase activity at Selvita S.A, Krakow, Poland.

---

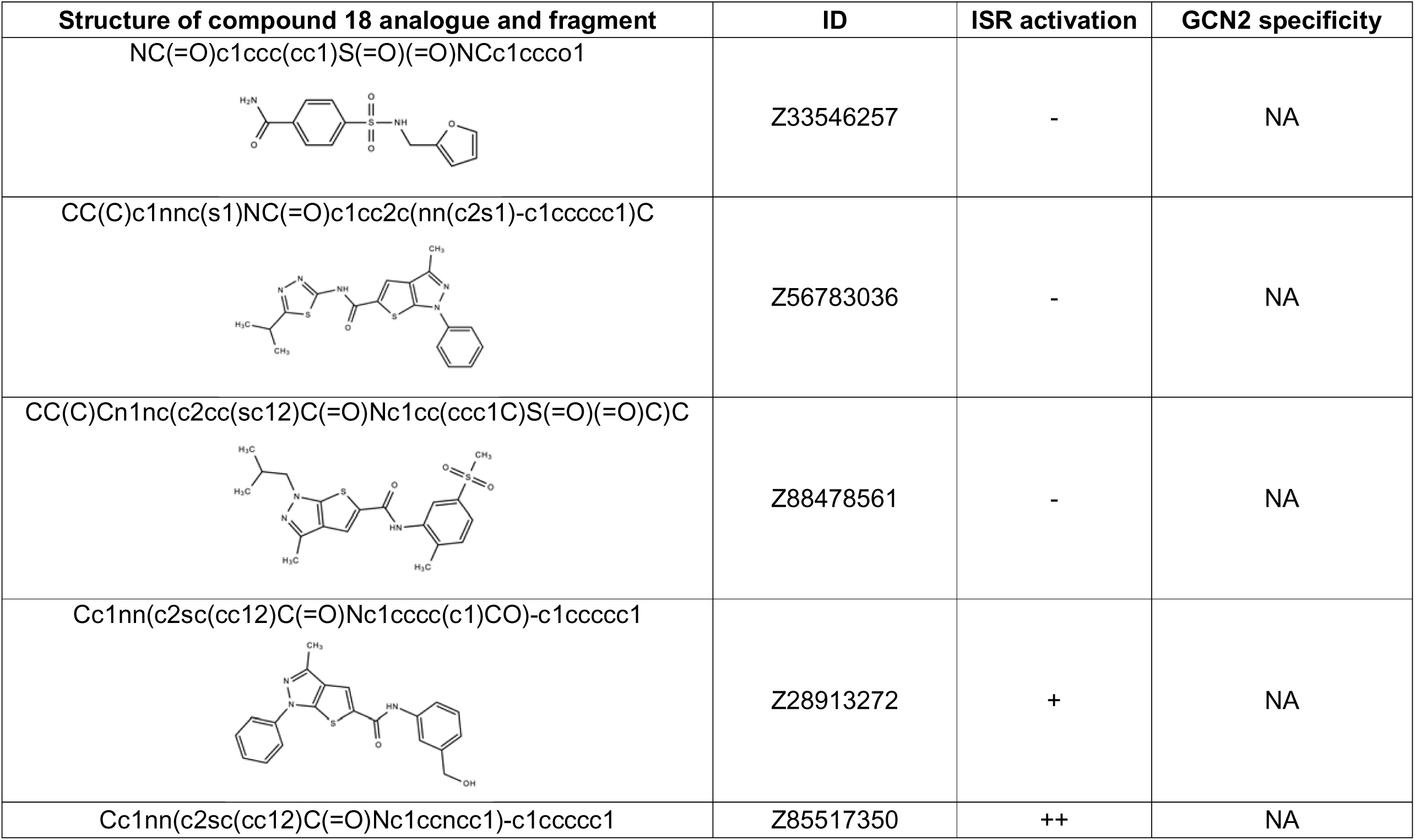
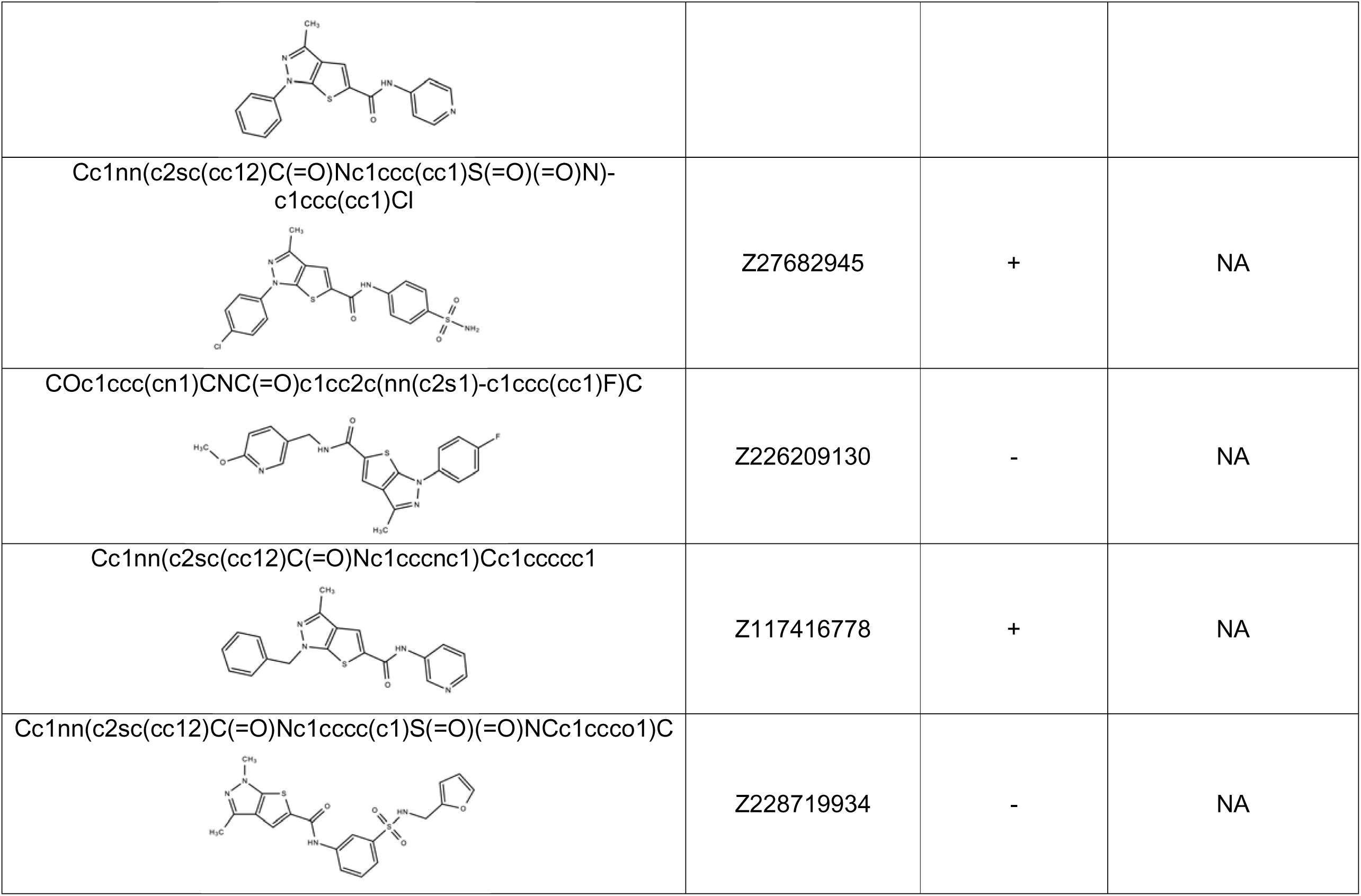
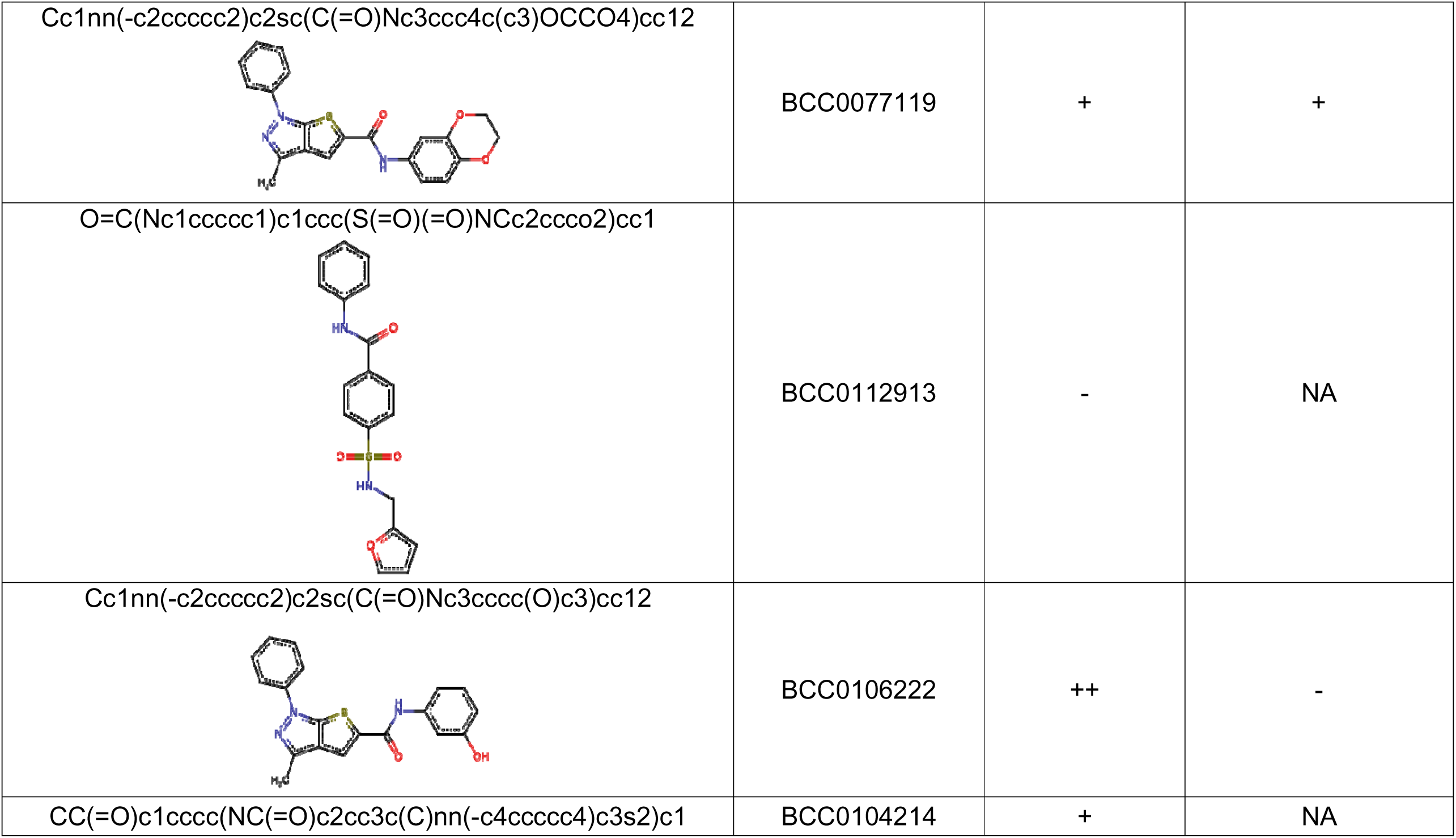
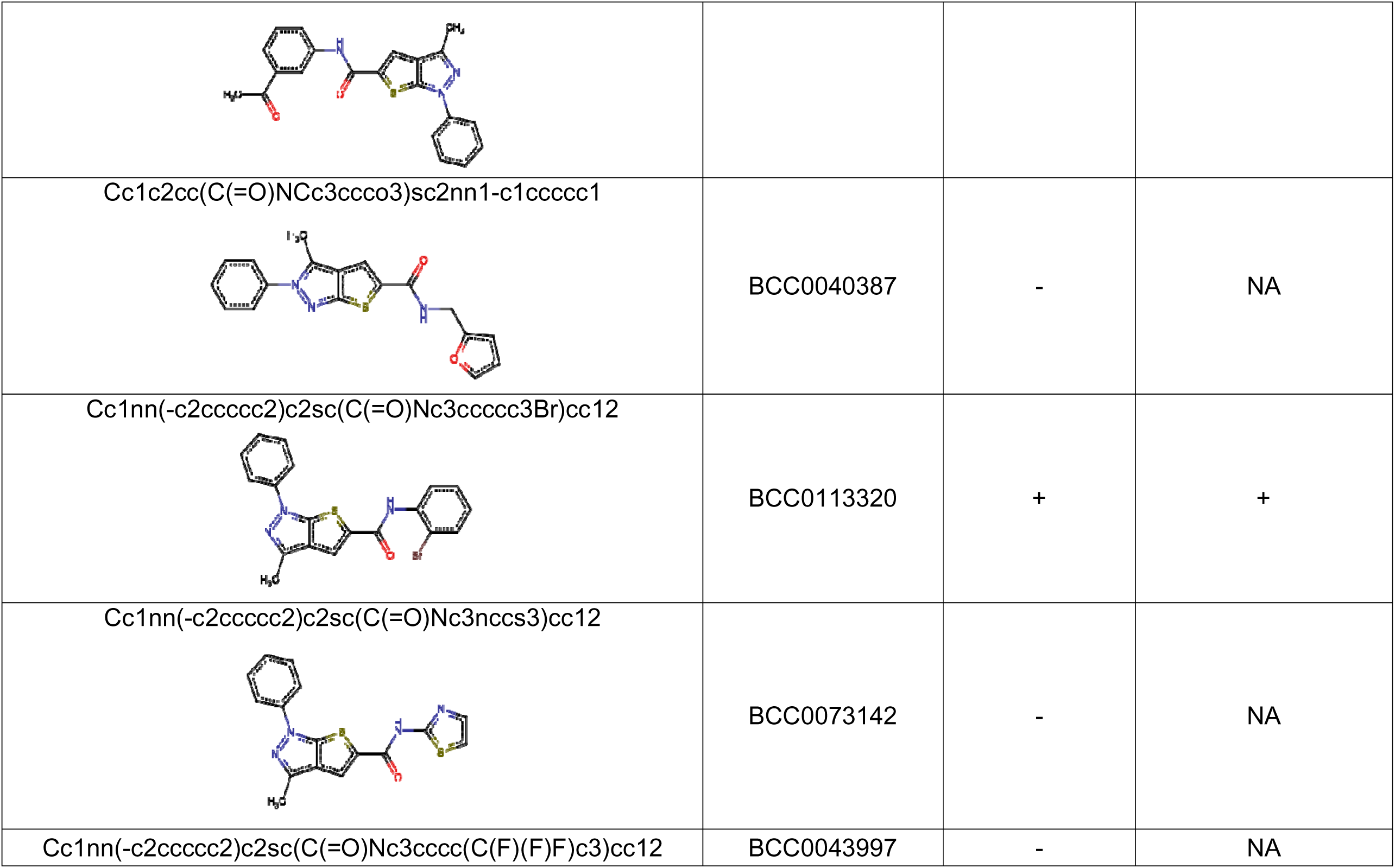
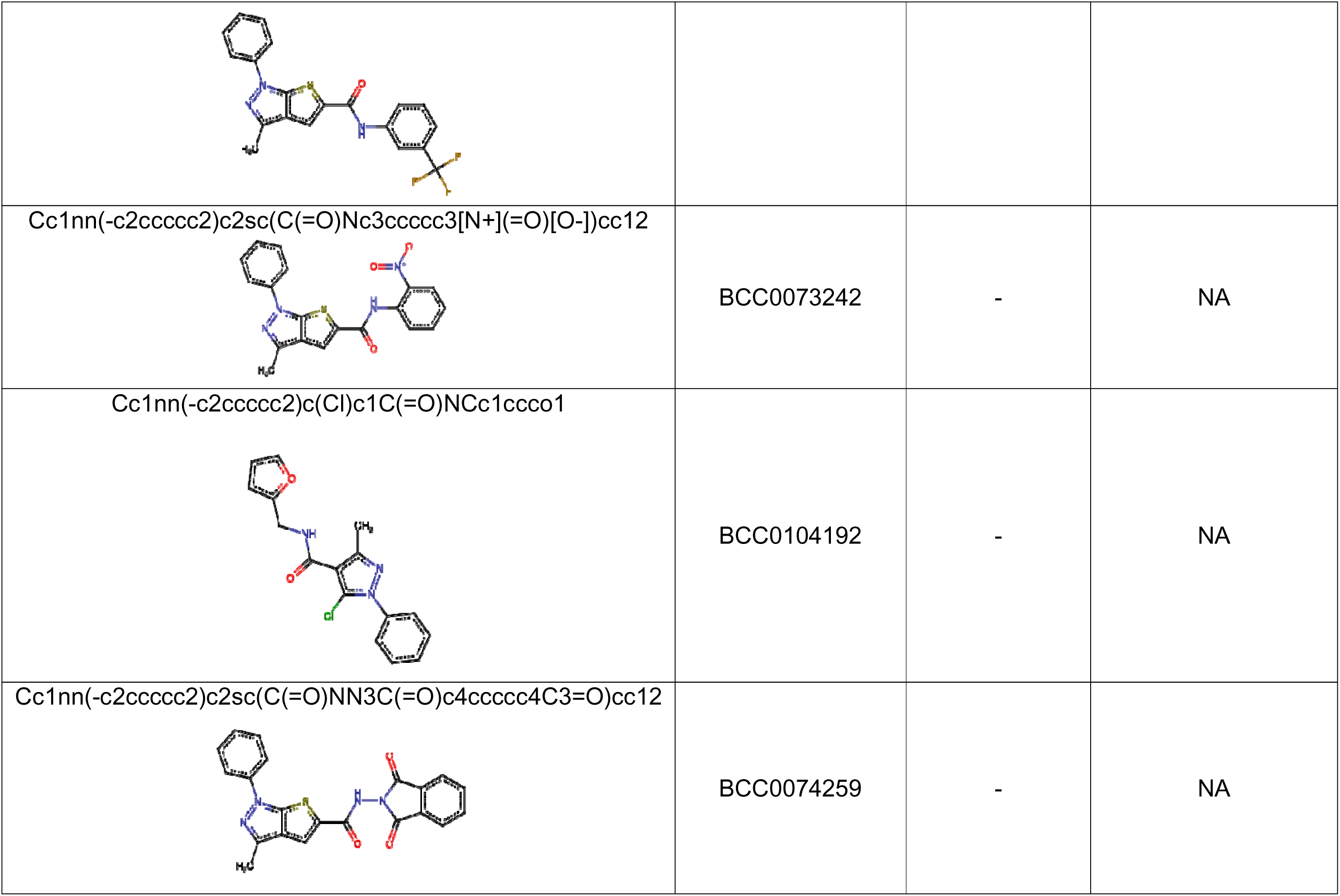
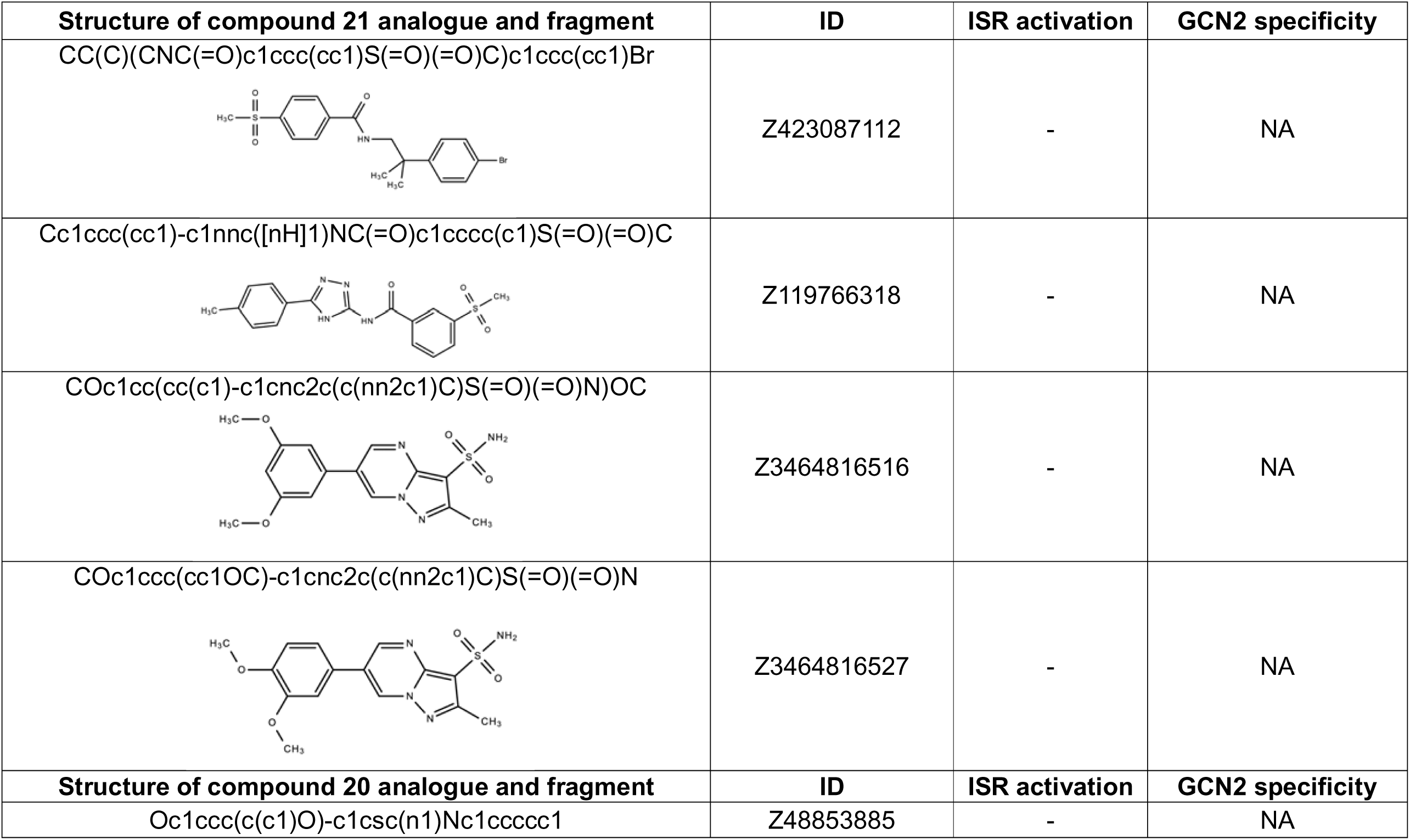
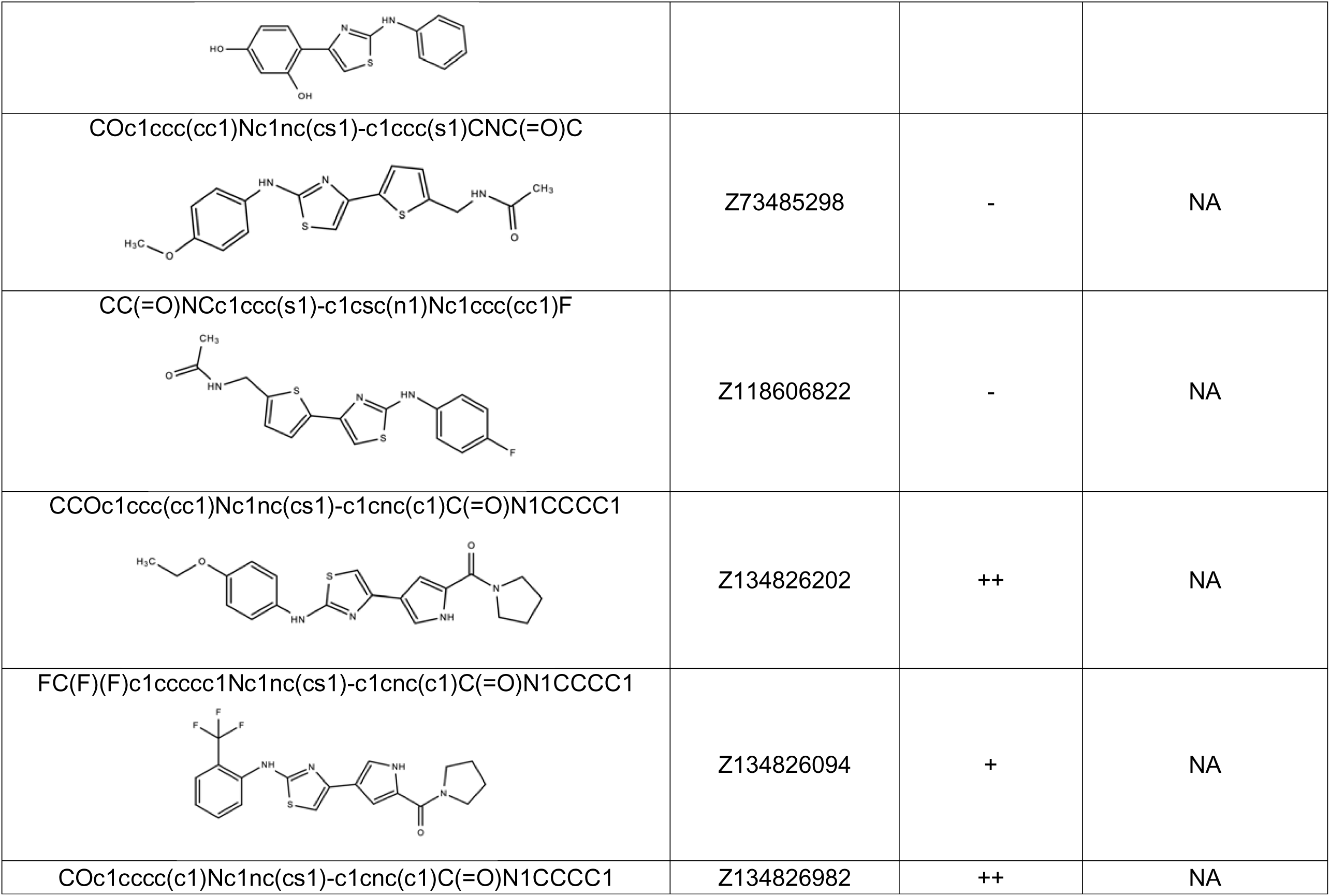
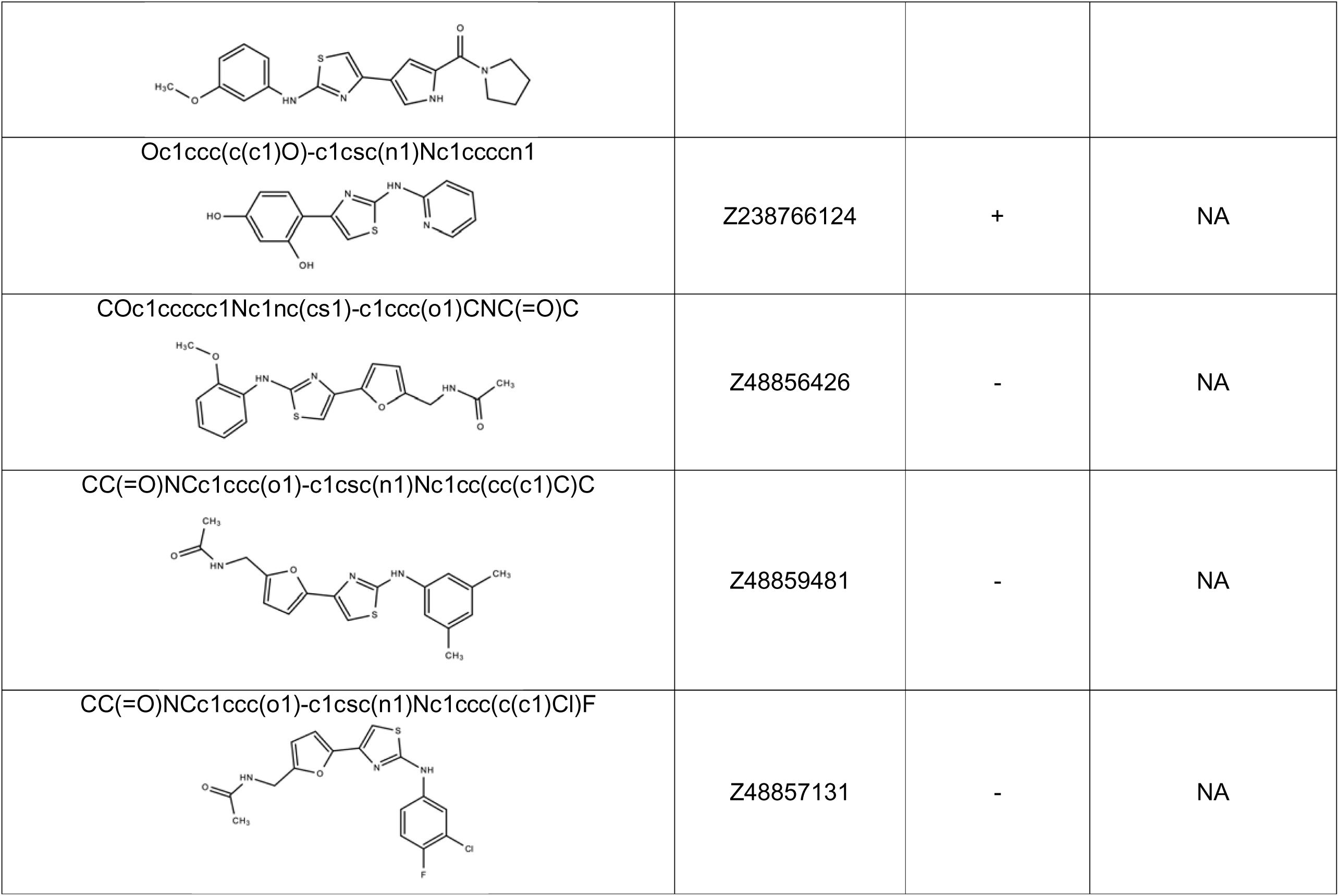
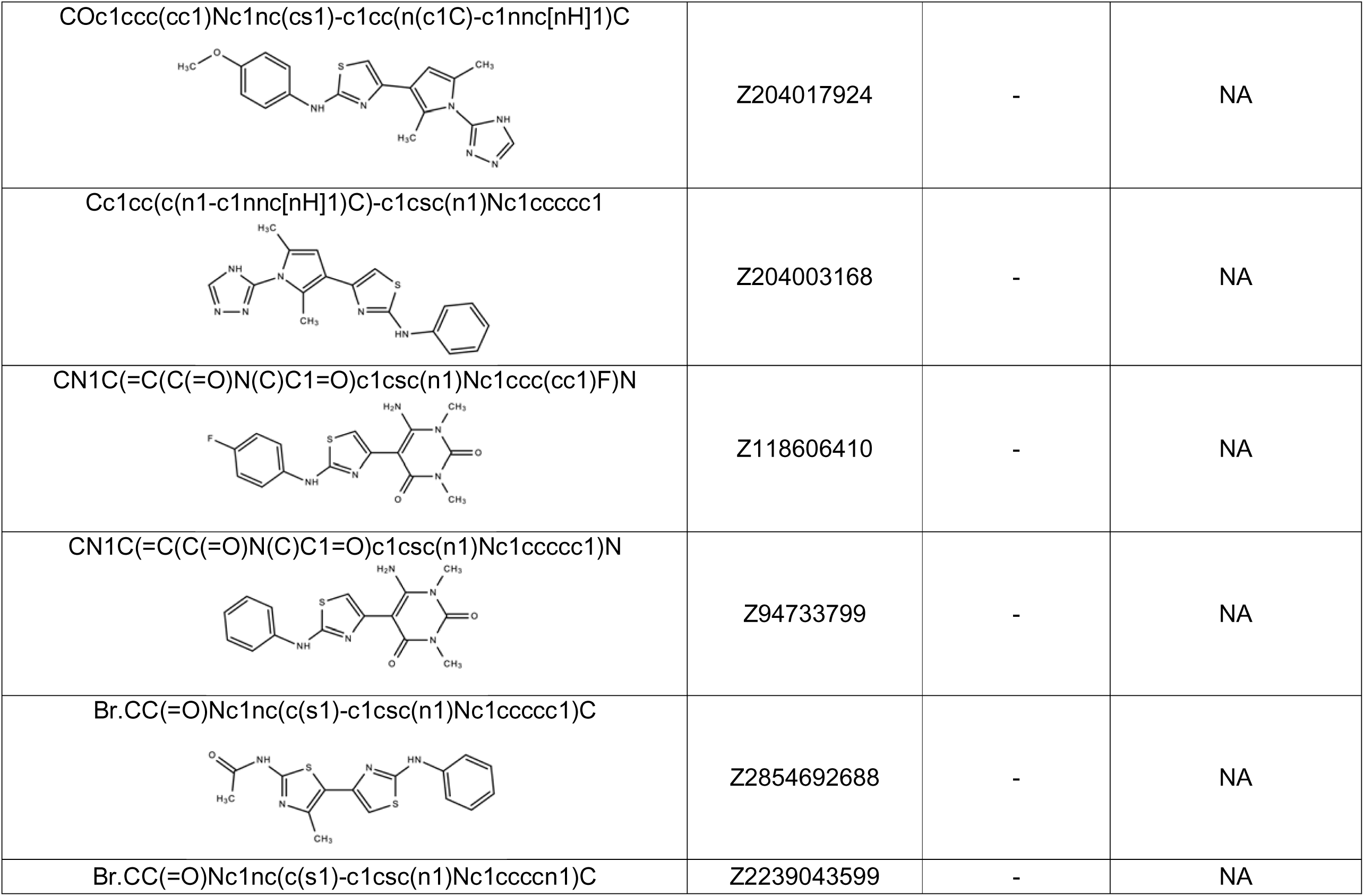
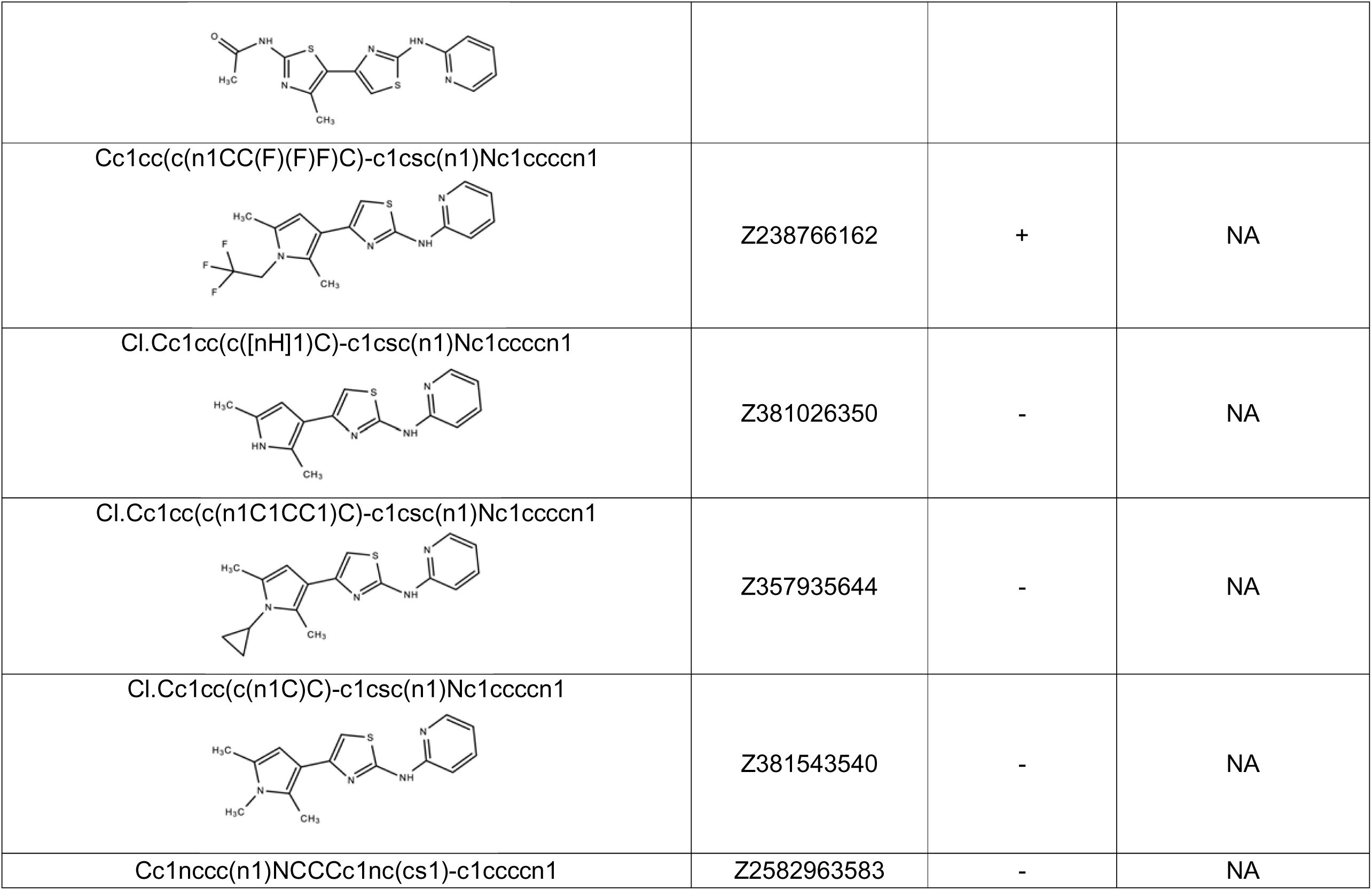
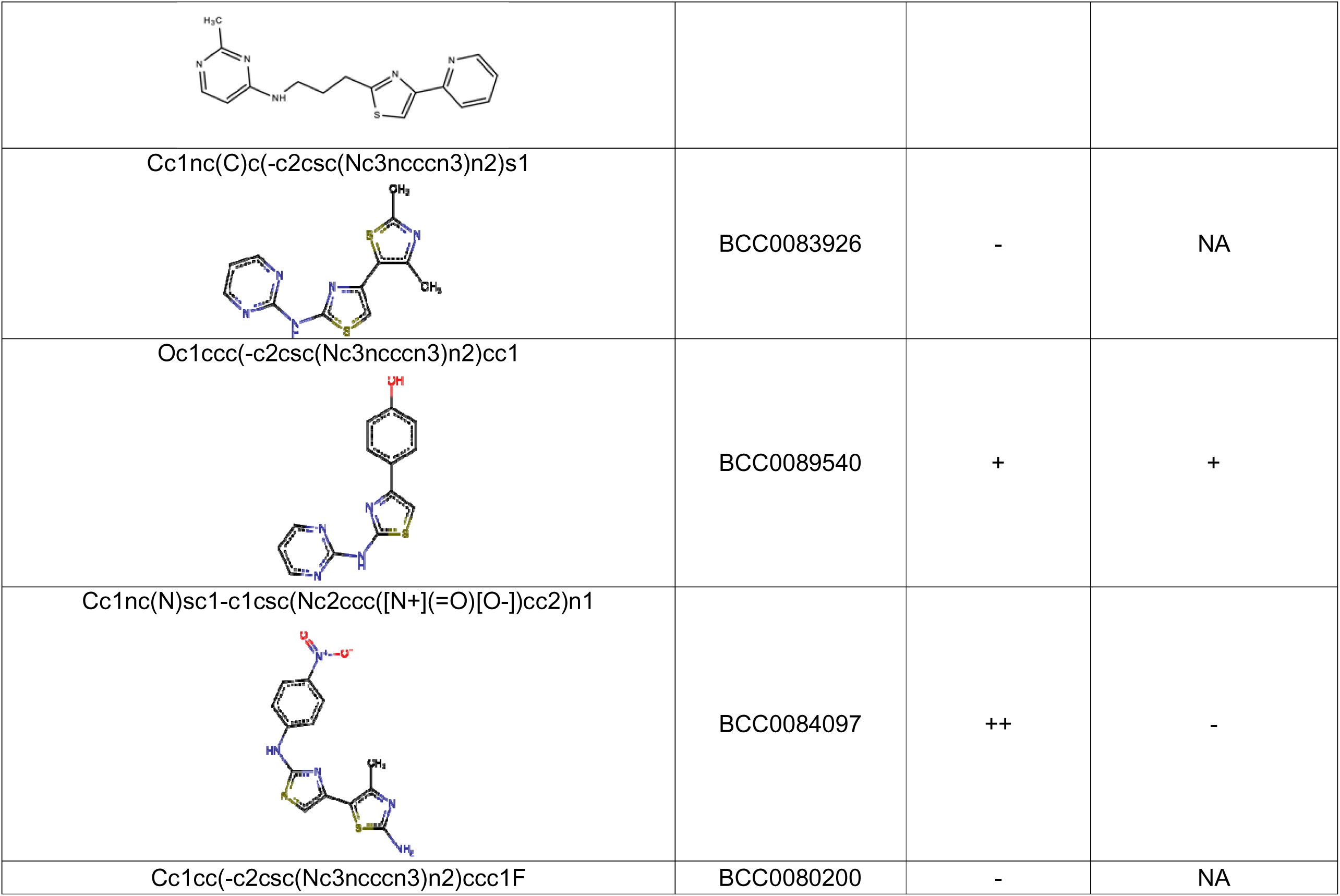
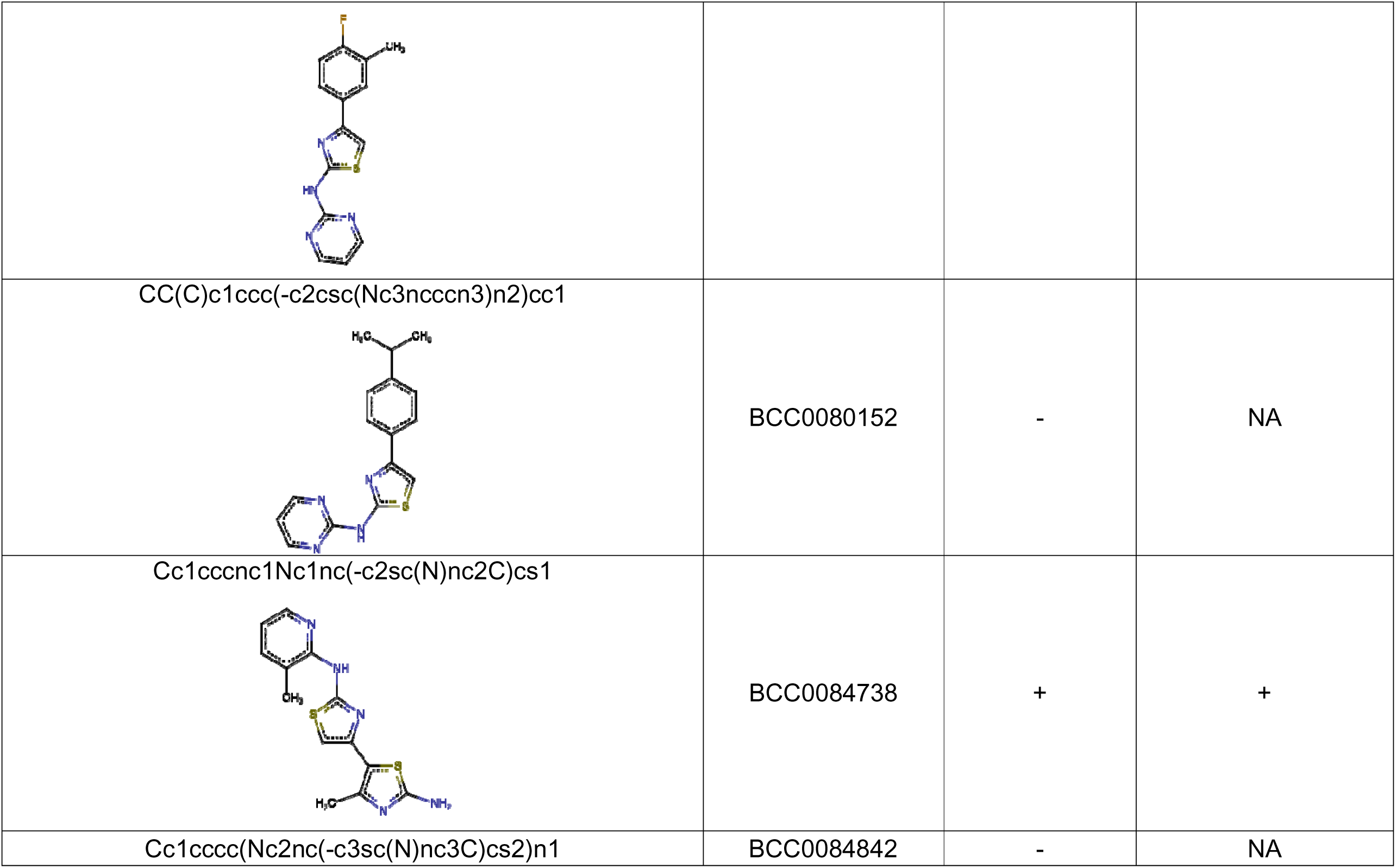
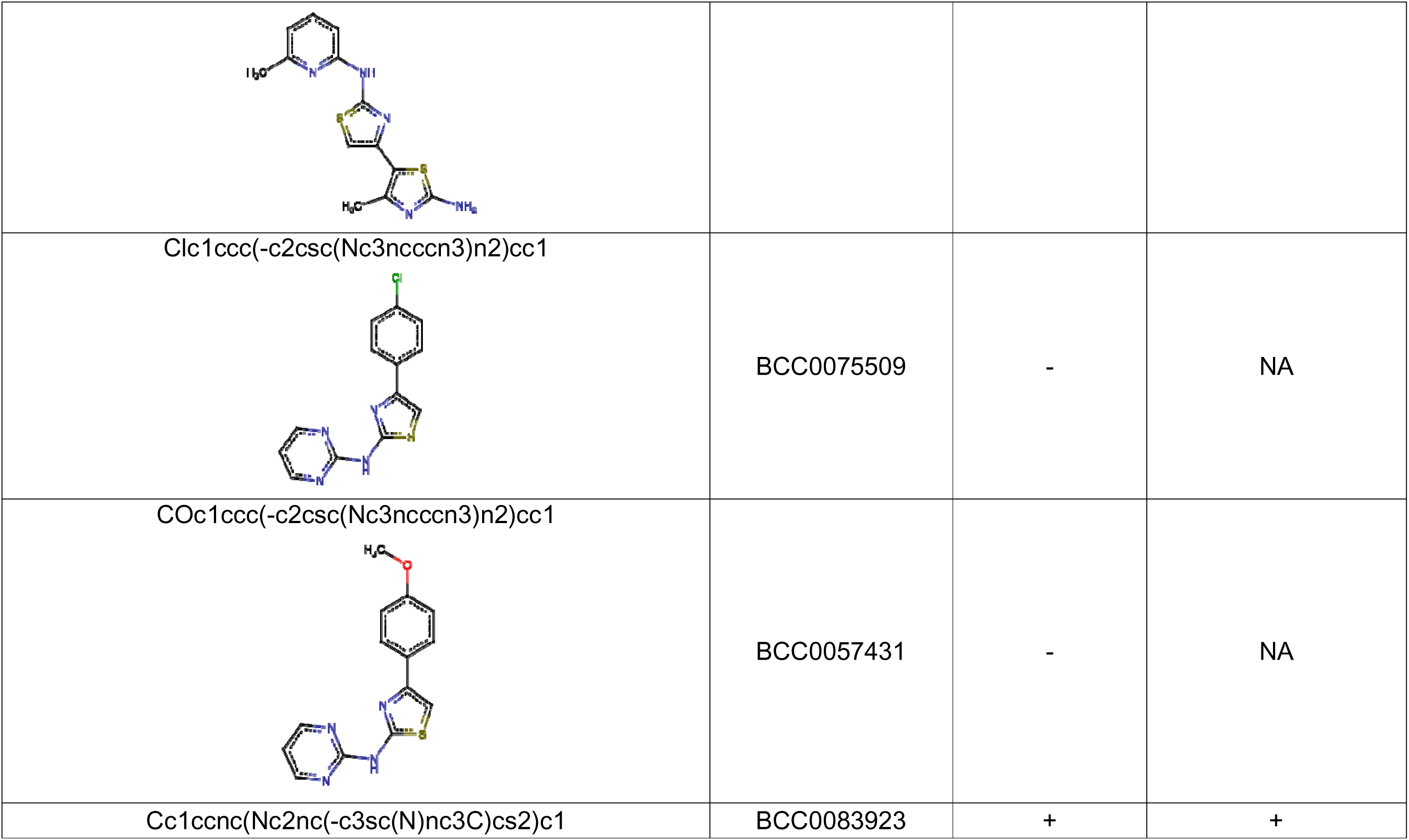
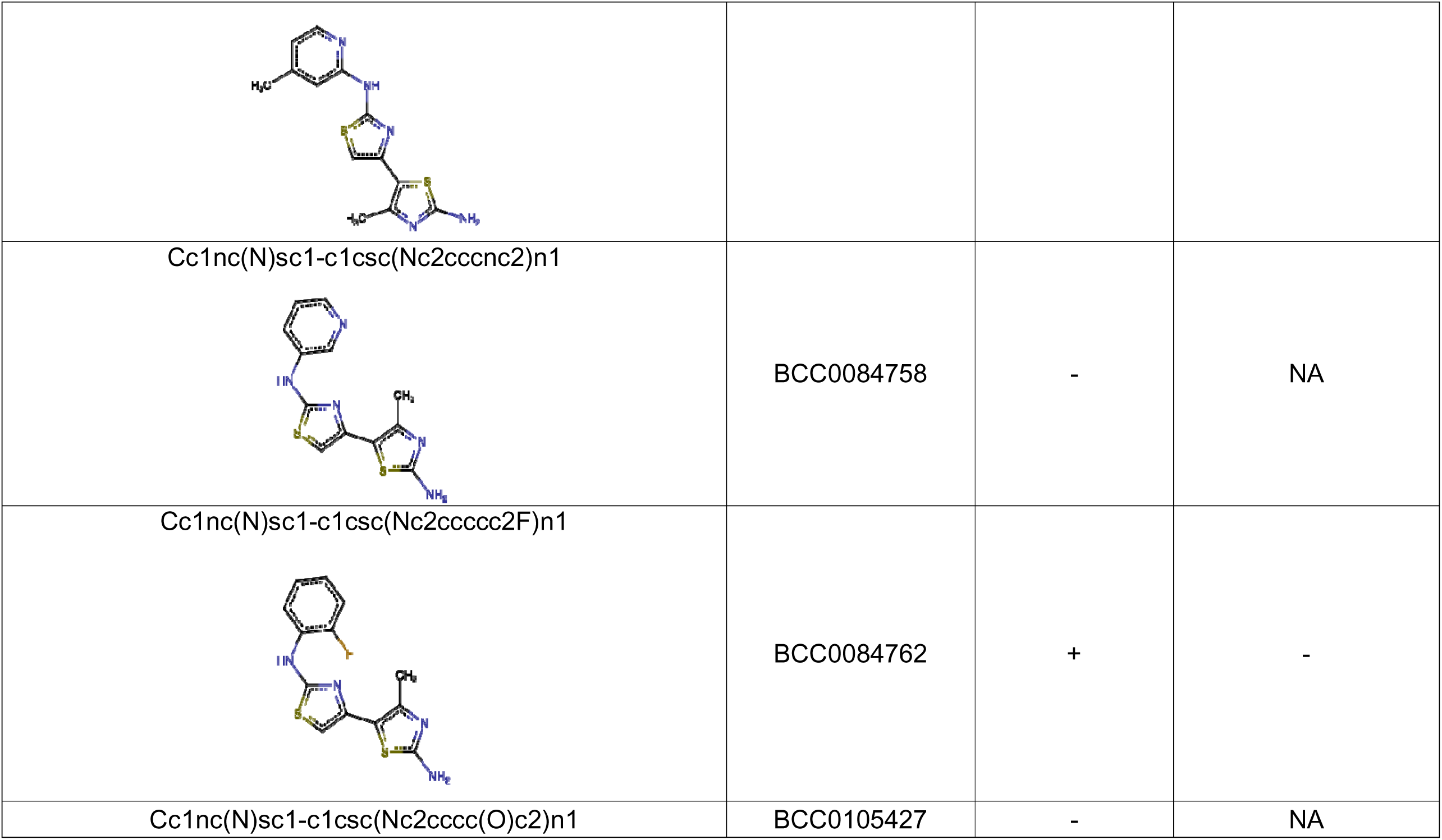
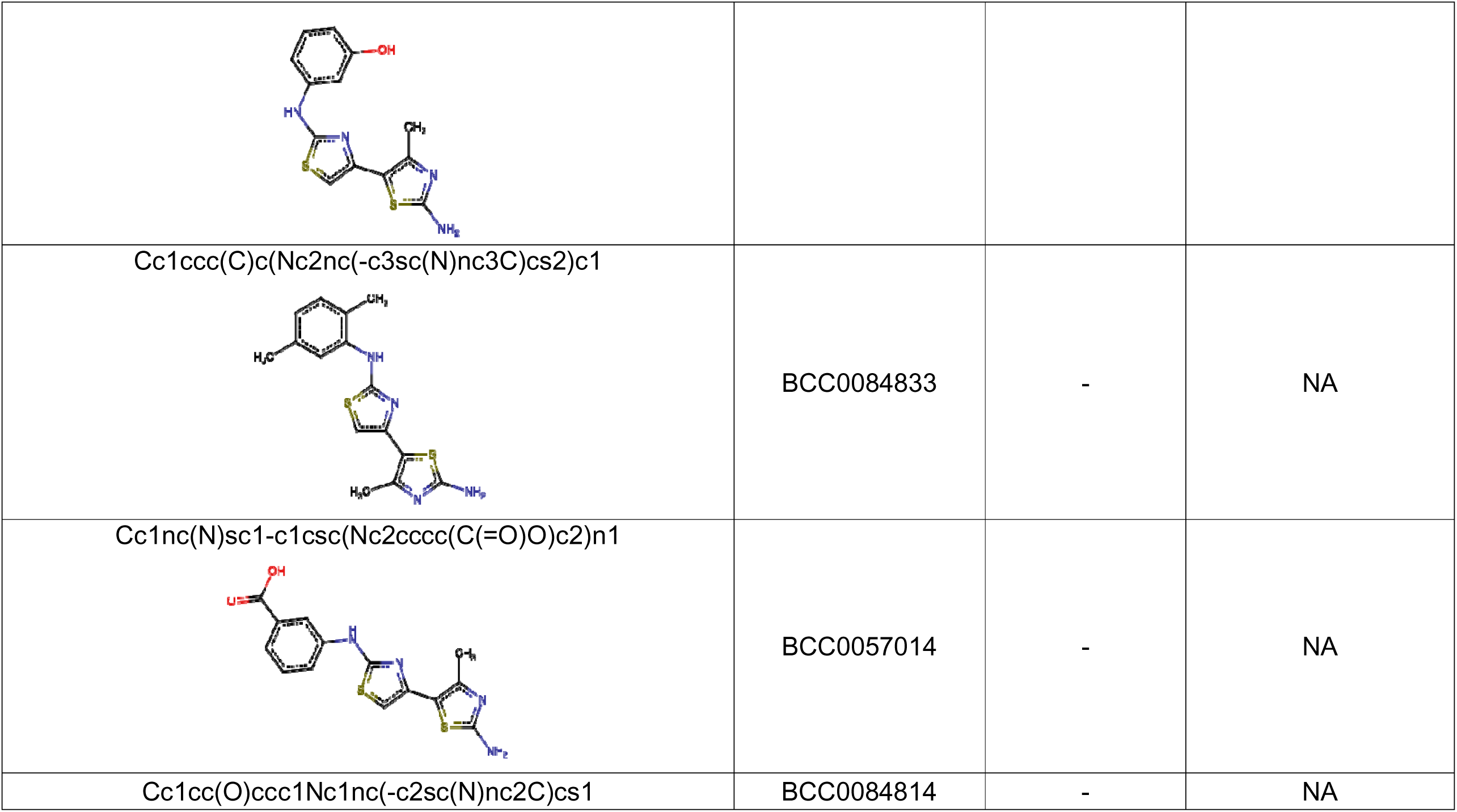
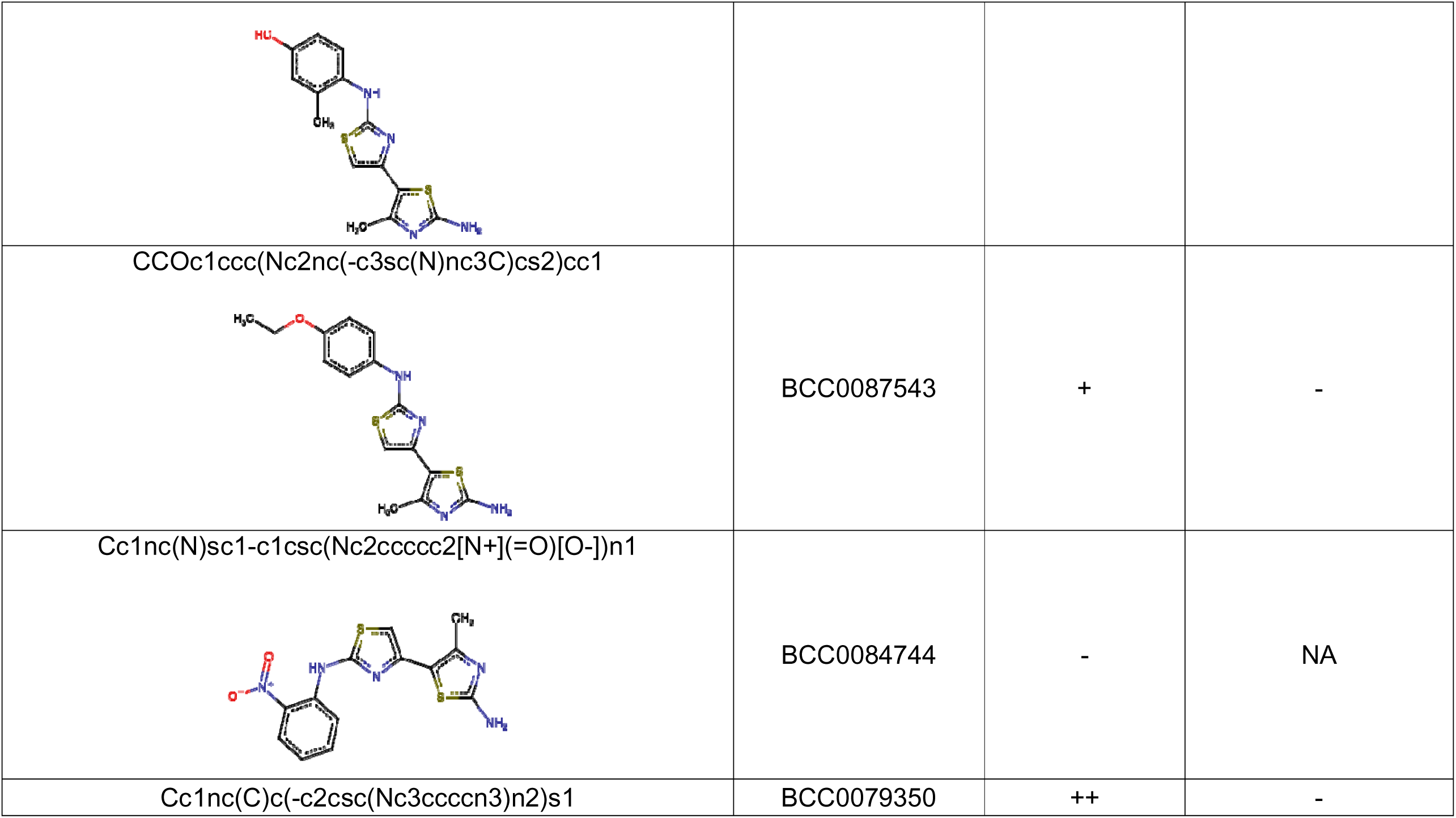
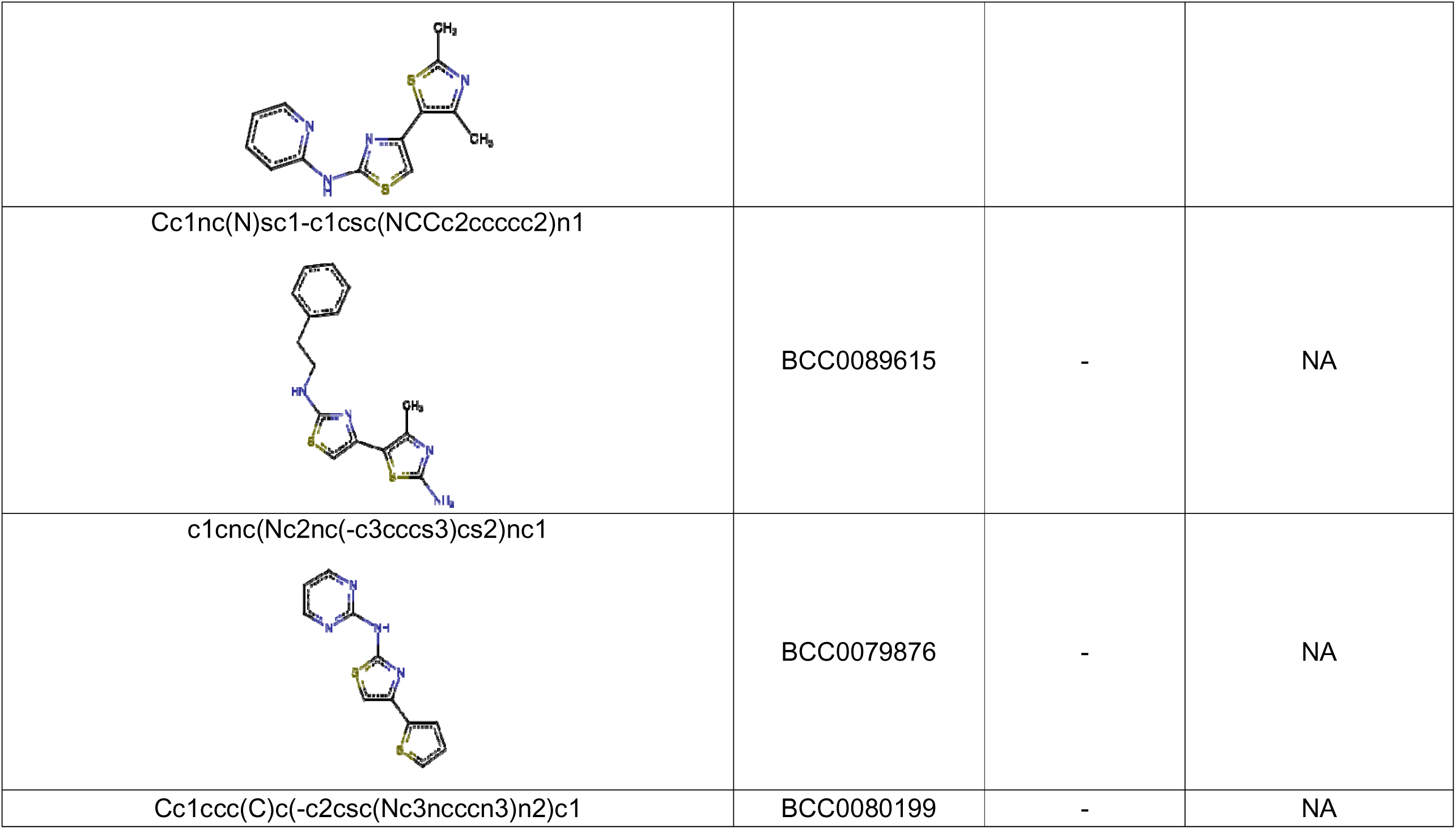
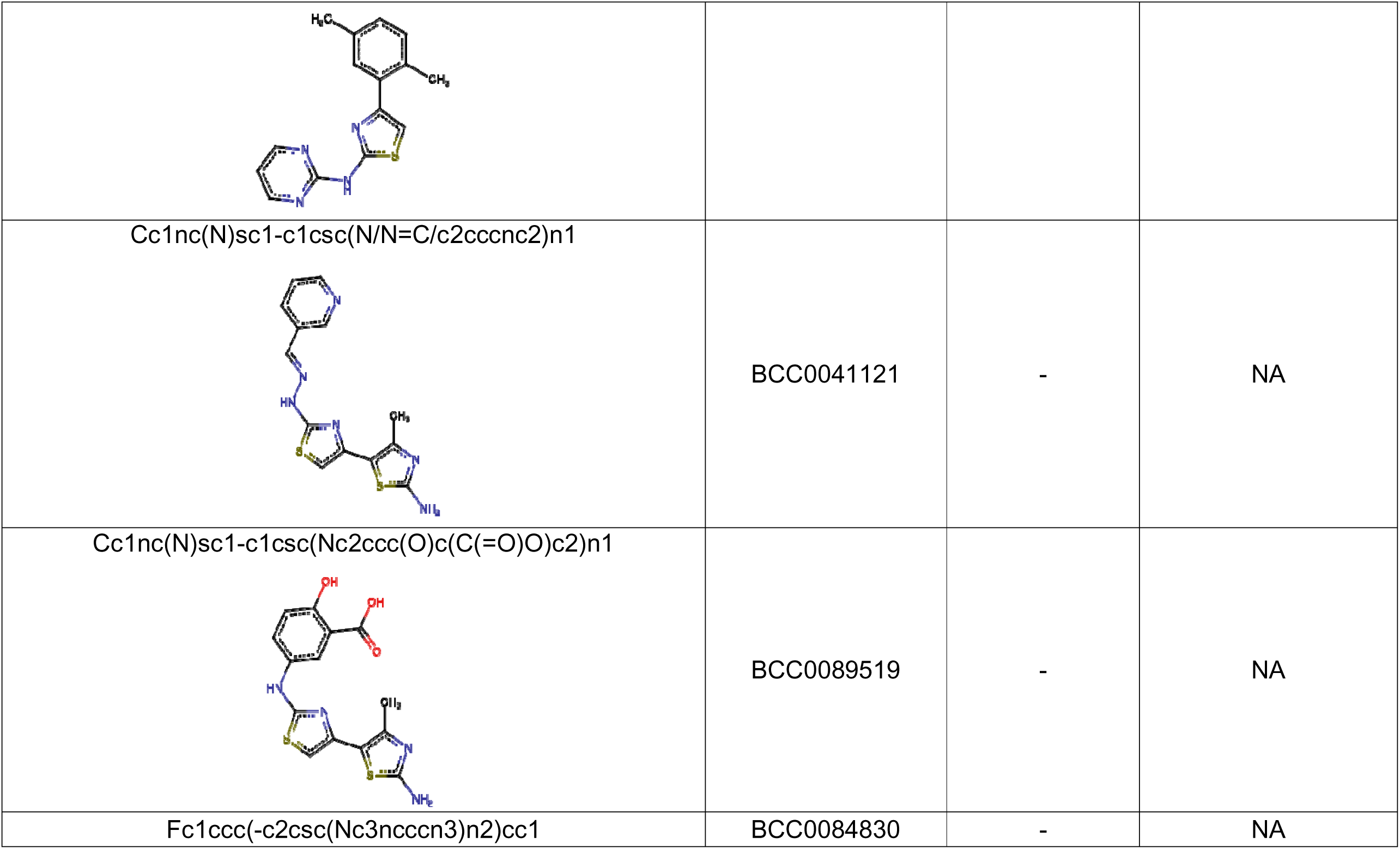
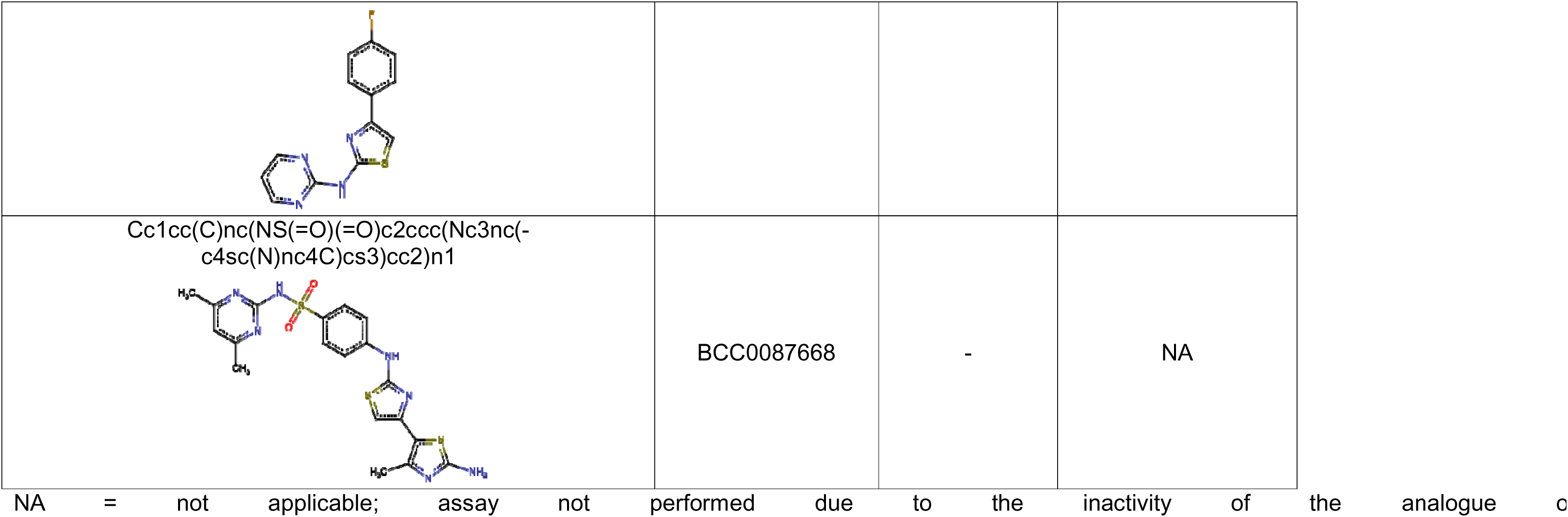

