## Supplementary figures and images for "A GCN1-independent activator of the kinase GCN2"

### Supplementary Figure 1

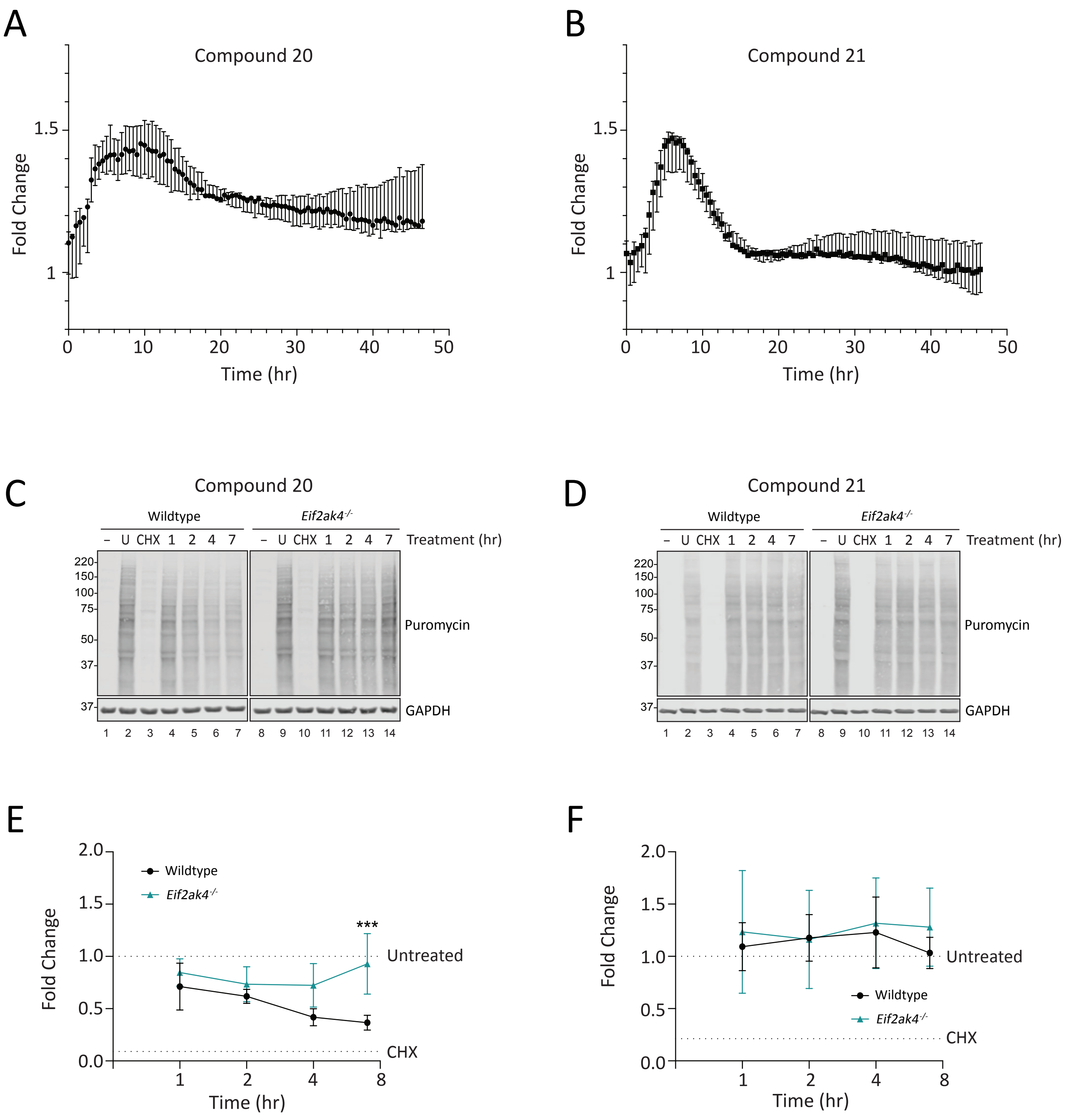

### Supplementary Figure 2

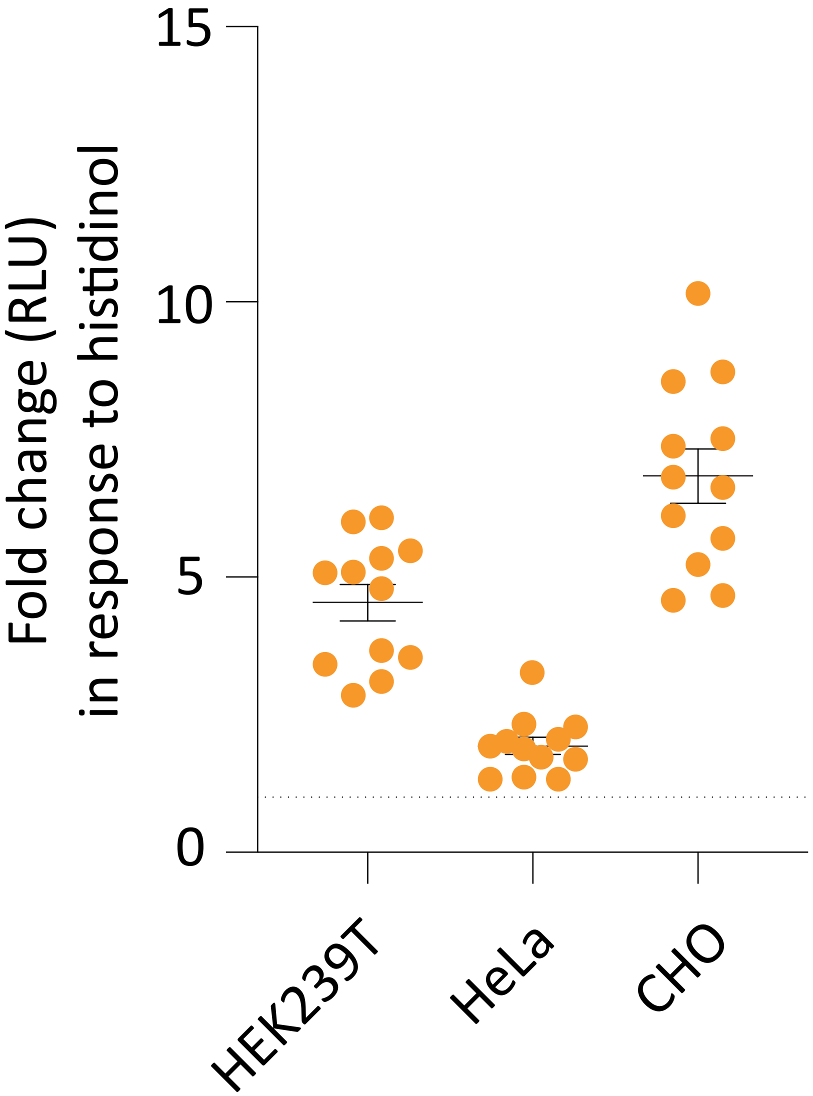

### Supplementary Figure 3

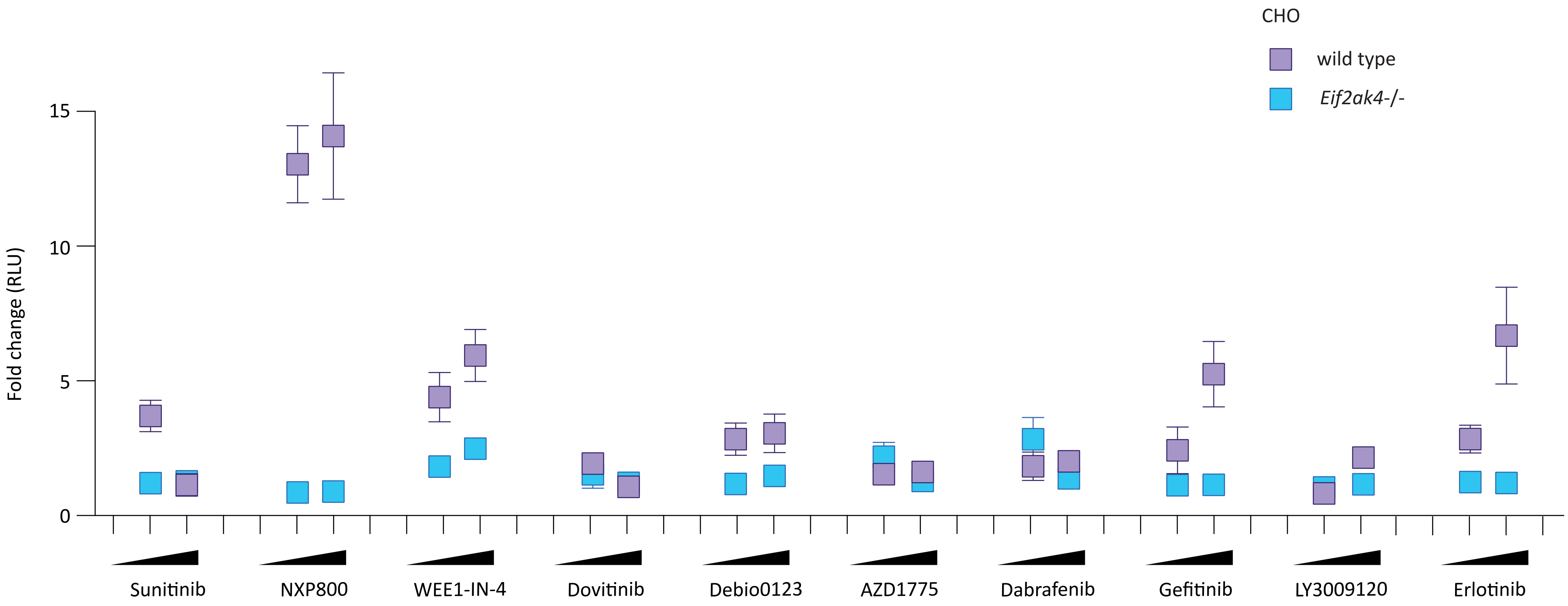
